# Allosteric Modulation of Pathological Ataxin-3 Aggregation: A Path to Spinocerebellar Ataxia Type-3 Therapies

**DOI:** 10.1101/2025.01.22.633970

**Authors:** Alexandra Silva, Sara Duarte-Silva, Pedro M. Martins, Beatriz Rodrigues, Débora Serrenho, Daniela Vilasboas-Campos, Andreia Teixeira-Castro, Julio Vieyto-Nuñez, Joel Mieres-Perez, Francisco Figueiredo, Joana Fraga, James Noble, Carter Lantz, Niki Sepanj, Daniela Monteiro-Fernandes, Sara Guerreiro, Andreia Neves-Carvalho, Joana Pereira-Sousa, Frank-Gerrit Klärner, Thomas Schrader, Joseph A. Loo, Annalisa Pastore, Elsa Sanchez-Garcia, Gal Bitan, Ana Luísa Carvalho, Patrícia Maciel, Sandra Macedo-Ribeiro

**Affiliations:** i3S-Institute for Research and Innovation in Health, Porto University, Porto, Portugal; Institute for Molecular and Cellular Biology (IBMC), Porto University, Porto, Portugal; Life and Health Sciences Research Institute (ICVS), School of Medicine, University of Minho, 4710-057 Braga, Portugal; ICVS/3B’s, PT Government Associate Laboratory, 4710-057 Braga, Portugal; Center for Neuroscience and Cell Biology (CNC-UC) & Center for Innovative Biomedicine and Biotechnology (CIBB), University of Coimbra, Coimbra, Portugal; Institute for Interdisciplinary Research, University of Coimbra, Coimbra, Portugal; Department of Biochemical and Chemical Engineering, TU Dortmund University, Dortmund, Germany; King’s College London, London, United Kingdom; Department of Chemistry and Biochemistry, University of California, Los Angeles, CA, USA; Department of Neurology, David Geffen School of Medicine, University of California, Los Angeles, Los Angeles, CA, USA; Faculty of Chemistry, University of Duisburg-Essen, Essen, Germany; Brain Research Institute and Molecular Biology Institute, University of California, Los Angeles, Los Angeles, CA, USA; Department of Life Sciences, University of Coimbra, Coimbra, Portugal

**Keywords:** Amyloid, Polyglutamine, Protein dynamics, Molecular therapies, Pre-clinical models

## Abstract

Spinocerebellar ataxia type 3 (SCA3) is a rare inherited neurodegenerative disease caused by the expansion of a polyglutamine repeat in the protease ataxin-3 (Atx3). Despite extensive knowledge of the downstream pathophysiology, no disease-modifying therapies are currently available to halt disease progression. The accumulation of protein inclusions enriched in the polyQ-expanded Atx3 in neurons suggests that inhibiting its self-assembly may yield targeted therapeutic approaches. Here it is shown that a supramolecular tweezer, CLR01, binds to a lysine residue on a positively charged surface patch of the Atx3 catalytic Josephin domain. At this site, the binding of CLR01 decreases the conformational fluctuations of the distal flexible hairpin. This results in reduced exposure of the nearby aggregation-prone region, which overlaps with the substrate ubiquitin binding site and primes Atx3 self-assembly, ultimately delaying Atx3 amyloid fibril formation and reducing the secondary nucleation rate, a process linked to fibril proliferation and toxicity. These effects translate into the reversal of synapse loss in a SCA3 cultured cortical neuron model, an improved locomotor function in a C. elegans SCA3 model, and a delay in disease onset, accompanied by reduced severity of motor symptoms in a SCA3 mouse model. This study provides critical insights into Atx3 self-assembly, revealing a novel allosteric site for designing CLR01-inspired therapies targeting pathological aggregation pathways while sparing essential functional sites. These findings emphasize that targeting allosteric sites in amyloid-forming proteins may offer unique opportunities to develop safe therapeutic strategies for various protein misfolding disorders.

## INTRODUCTION

Spinocerebellar Ataxia type 3 (SCA3), also known as Machado-Joseph disease (MJD), is a rare, neurodegenerative, hereditary ataxia with no disease-modifying treatments. SCA3 results from the expansion of a trinucleotide CAG repeat in the ATXN3 gene, which is translated into an expanded polyglutamine (polyQ) tract in the ataxin-3 protein (Atx3). Atx3 is a modular deubiquitinating enzyme displaying high conformational plasticity, which is composed of a globular catalytic Josephin domain (JD) and a flexible C-terminal tail containing the polyQ tract and two or three ubiquitin-interacting motifs (UIMs).[1] Aggregates containing Atx3 with pathological polyQ tract lengths (>61Q) represent a key disease fingerprint and accumulate in degenerating brain regions, including the cerebellum, brain stem, substantia nigra, and striatum, as well as the spinal cord.[2–4] The exact cause of neurotoxicity is still debated, but several cell and animal disease models show that abnormal aggregation of pathological Atx3 contributes to the underlying neuronal damage in SCA3[2, 5–9] and other polyQ disorders.[9–15] As a result, tackling aberrant Atx3 self-assembly is one of the main approaches pursued to find disease-modifying therapies for SCA3 and other polyQ diseases.[16–19]

Biophysical studies have shown that both wild-type and pathological forms of Atx3 can assemble into amyloid-like structures in vitro.[20–22] An aggregation-prone region located in the JD (^73^GFFSIQVISNALKVWGLELILFNS^96^), overlapping with the substrate ubiquitin binding site, plays an important role in initiating the Atx3 self-assembly process.[20–22] However, only the expansion of the polyQ beyond the pathological threshold triggers the formation of mature, SDS-resistant fibrils.[23–25] Studies using non-pathological Atx3 demonstrated the occurrence of transient populations of soluble oligomers in parallel with the thermodynamically favored amyloid pathway.[26, 27] Given the complexity of the Atx3 self-assembly process and the heterogeneity of its assembly states, which include monomers, dimers, amyloidogenic and non-amyloidogenic oligomers, amyloid protofibrils, and bundled amyloid fibrils, targeting Atx3 self-assembly is challenging. Therefore, to develop effective inhibitors that target neurotoxic Atx3 aggregation, dissecting their molecular mechanism of action and comprehending how they interfere with the various steps involved in Atx3 self-assembly is crucial.

Here, we investigated the molecular tweezer CLR01 (Fig. 1a), a Janus-type supramolecular ligand using a distinctive recognition mode, which acts as a broad-spectrum inhibitor of abnormal protein self-assembly.[28] CLR01 possesses an electron-rich cavity and adjacent phosphate anions that can accommodate a lysine or an arginine side chain, and a convex exterior capable of docking onto hydrophobic clefts on the target proteins[29]. By combining in silico, in vitro, and in vivo studies, we evaluated the impact of CLR01 on Atx3 self-assembly and investigated its effect on disease-related phenotypes in cell and animal models of SCA3. Our results show that CLR01 binds to a surface patch close to the termini of the JD domain of Atx3, acting as an allosteric modulator that restrains the conformational fluctuations required to expose the distal aggregation-prone segment, which is part of the Atx3 amyloid fibril core[25]. CLR01 effectively remodeled the Atx3 aggregation pathway and not only reduced the rate of secondary nucleation, a crucial step for the proliferation of amyloid fibrils, but also disrupted pre-formed fibrils. In addition, CLR01 reverted synaptic defects in primary neurons expressing polyQ-expanded Atx3 and reduced motor deficits in a SCA3 *C. elegans* model. To further validate its efficacy against Atx3, we tested CLR01 using the CMVMJD135 transgenic mouse model of SCA3, which exhibits quantifiable motor symptoms similar to those observed in patients with SCA3.[6] Chronic and early symptomatic treatment with CLR01 demonstrated significant therapeutic benefits, delaying symptom onset and improving motor function in correlation with rescued motor-neuron pathology. These outcomes underscore the potential of this newly identified CLR01 binding site as a valuable target in the development of innovative lead molecules to address the toxicity linked to pathological Atx3 aggregation in SCA3.

**Figure 1.**
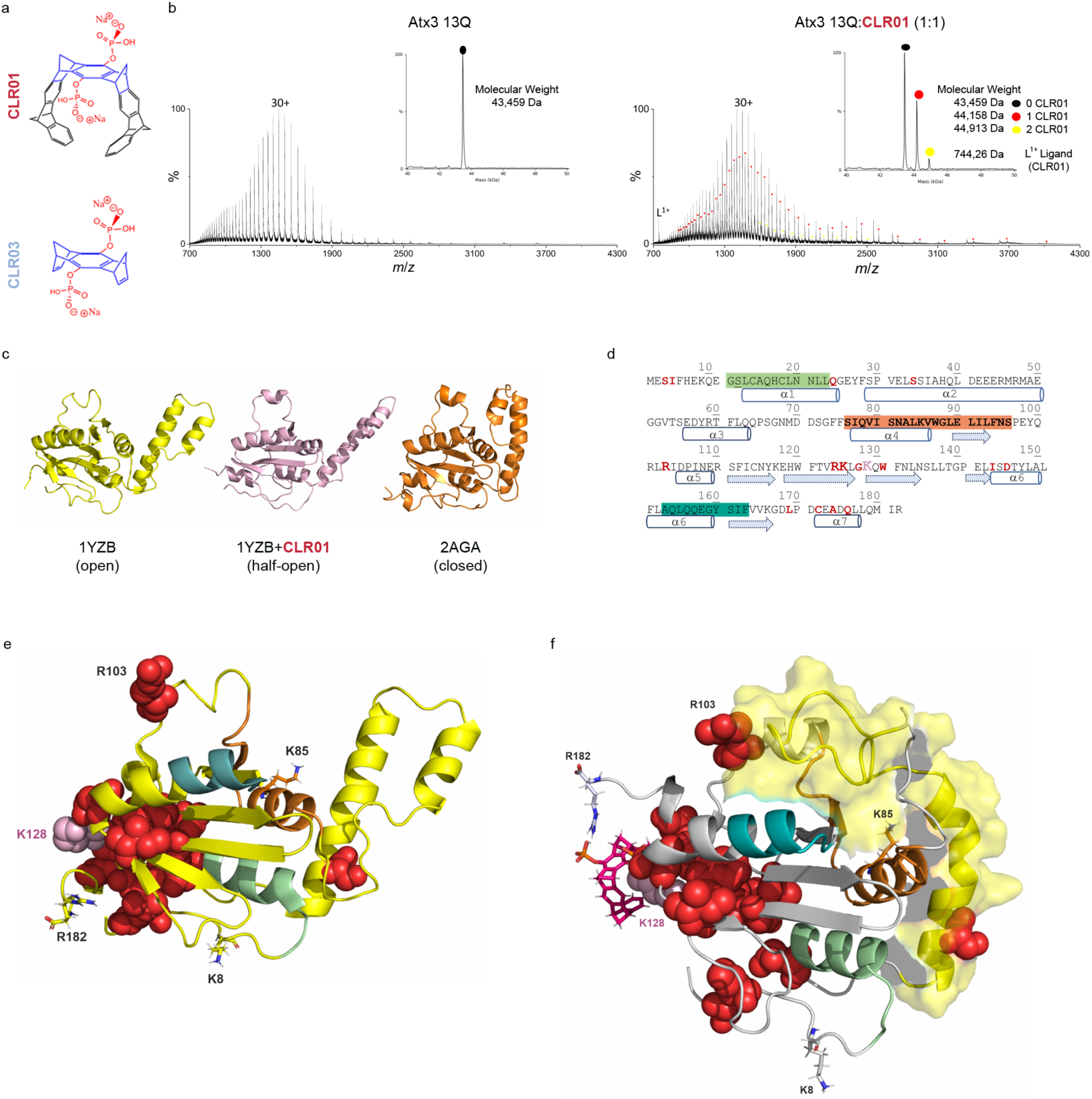
CLR01 interacts with Atx3 and induces conformational changes in the JD. **a)** Chemical structure of the molecular tweezer CLR01 and the control molecule CLR03. **b)** ESI-MS spectra of wild-type Atx3 (Atx3 13Q) and CLR01 at a 1:1 molar ratio. The inset shows the deconvoluted spectra and highlights the two CLR01 binding events. **c)** Cartoon representation of three states of JD: Experimental NMR models of the JD in open (PDB ID: 1YZB,[33] yellow) and closed (PDB ID: 2AGA,[34] orange) conformations. A representative structure (light pink) of the “half-open” conformation of the JD is shown (most populated cluster from the GaMD simulations in complex with CLR01). **d)** Amino acid sequence of JD with residues undergoing chemical shifts in the NMR HSQC spectra upon CLR01 binding in red. Secondary structure elements in the JD are shown below the amino acid sequence with helix α1 in green, the aggregation-prone segment (α4-β1) in bold and orange, and helix α6 in teal. **e)** Cartoon representation of the JD open conformation with the residues affected by CSP upon addition of CLR01 shown as red spheres. K128, a key residue in the interaction between CLR01 and JD, is highlighted as pink spheres. Other regions are colored as in panel d). **f)** Cartoon representation of the third most populated cluster from the GaMD simulations of JD (open conformation) with a single CLR01 molecule (magenta sticks) bound at K128 (pink spheres). Residues displaying a chemical shift in the HSQC spectrum are highlighted as red spheres; CLR01 is represented as dark pink sticks; R182 and other positively charged residues, corresponding to predicted CLR01 binding sites according to GaMD simulations and Gibbs binding energy estimations, are shown as sticks. This structure highlights a state where the hairpin (yellow surface) moves toward the aggregation-prone region (orange).

## RESULTS

### CLR01 binds to Atx3, inducing conformational changes in the helical hairpin

Several studies demonstrated that JD dimerization primes Atx3 self-assembly[20, 22, 25] and that fibril formation is critically dependent on the exposure of an amyloidogenic region that includes an exposed lysine residue. [21, 24, 25] As CLR01 (**Figure 1a**) can bind to exposed lysine or arginine residues, we hypothesized it may disturb the electrostatic and hydrophobic interactions within the aggregation-prone region, thereby interfering with Atx3 aggregation.

To evaluate CLR01’s interaction(s) with Atx3 and gain insights into its impact on Atx3 conformational dynamics, we conducted native electrospray ionization mass spectrometry (ESI-MS) studies, alongside Gaussian accelerated molecular dynamics (GaMD)[30] simulations. The addition of CLR01 at a 1:1 ratio to wild-type Atx3 (Atx3 13Q) revealed the formation of Atx3:CLR01 complexes with a K_D_=8 µM, which appeared as distinct distributions with shifts to higher m/z values for each charge state (**Figure 1b**). Upon deconvolution of the mass spectra, two binding events were observed, reflecting the formation of 1:1 and 1:2 Atx3:CLR01 complexes. Additionally, we observed that similar complexes were formed when CLR01 was added to the isolated JD (K_D_ 12 µM) or to a disease-related variant in which the polyQ tract was expanded to 77Q (Atx3 77Q, K_D_ 14 µM) (**Supporting Information Figure S1a, b**). The formation of CLR01-bound complexes with the isolated JD indicates that CLR01 molecules bind to this domain despite several potentially highly exposed K/R residues in the Atx3 C-terminal tail. Ion mobility coupled to mass spectrometry experiments indicated that wild-type and pathological Atx3, as well as the isolated JD, bound to CLR01, shifted toward shorter drift times, i.e., formed more compact conformers (**Supporting Information Figure S1c, d**).

To further understand the impact of CLR01 binding on Atx3 conformational fluctuations, we performed GaMD[30] simulations. Without CLR01, JD (PDB ID: 1YZB) had an open conformation, with the helical α2-α3 hairpin distant from the catalytic subdomain.[31, 32] CLR01 molecules were placed around the surface accessible lysine and arginine residues, resulting in a 1:9 CLR01 complex (**Supporting Information Figure S2**). In the presence of CLR01, the system evolved to a “half-open” conformation, representing an intermediate state between the experimentally reported “open” (PDB ID: 1YZB) and “closed” (PDB ID: 2AGA) conformations (**Figure 1c**, **Supporting Information Figure S3a-d**). Therefore, we also performed analogous GaMD simulations with the closed state (PDB ID: 2AGA) as the starting geometry. Based on our simulations, it is evident that in the absence of CLR01, the JD is better represented by the open conformation (PDB ID: 1YZB), in agreement with previous studies.[31, 33] Additionally, the JD in the closed conformation also transitions to a “half-open” configuration in the presence of CLR01 (**Supporting Information Figure S3d**). As a result, the open conformation was selected for further computational studies. The analysis of the root-mean-square-fluctuation (RMSF) for each residue over the complete GaMD trajectories confirmed that the largest conformational variability is located on the hairpin region of the JD (**Supporting Information Figure S3**).

Taken together, we found that CLR01 binds to JD and modulates the distribution of Atx3 conformers towards intermediate structures between the open and closed states, likely mediated by changes in the mobility of the JD helical hairpin.

### The primary site of CLR01 binding on JD is K128

To finely map the CLR01 binding sites on Atx3, we combined nuclear magnetic resonance (NMR) analysis with GaMD simulations and Gibbs energy estimations. We specifically focused on the effects of CLR01 on JD because this domain is known for driving the early stages of Atx3 aggregation[22, 24, 25, 35] and binds CLR01 (**Figure 1b, Supporting Information Figure S1a**). Enhanced sampling simulations of the JD in complex with nine CLR01 molecules bound to the R/K residues that were sterically accessible (**Supporting Information Figures S2, S4a)** revealed a higher affinity for lysine residues compared to arginines (**Supporting Information Figure S4b**). Further, we estimated binding Gibbs energy changes for the complexes between CLR01 and selected residues of JD using the CL-FEP approach.[36] The Gibbs energy calculations reveal seven sites where binding of CLR01 to exposed lysine and arginine residues is favored (**Supporting Information Figure S4c**). NMR titration of JD with CLR01 showed that the tweezer induced several clear chemical-shift-perturbation (CSP) effects on specific resonances. The residues most affected, indicated by the total disappearance of the amide proton resonance, are G127, K128, I143, and A174. The resonances of S3, I4, Q24, S34, R103, R124, K125, W130, L169, C172, and Q176 are perturbed and shifted (**Figure 1d, Supporting Information FigureS4d**). The main binding site is centered on the positively charged residue K128, which is located in the β2-β3 loop, next to helix α7 (^174^ADQ^176^), the N-terminal residues ^2^ESI^4^, and the C-terminal portion of strand β5 (I143). Although distant in the sequence, most of the residues are clustered in space in the same area, which contains the N- and C-termini of the JD. The CLR01 interaction site includes an exposed hydrophobic patch formed by the aliphatic moieties of Q176 and L177 side chain, which likely establish additional hydrophobic interactions with K128-bound CLR01. Binding to CLR01 is in agreement with the estimated Gibbs energy changes, showing that the binding of CLR01 to the exposed K128 residue on the JD was thermodynamically favored (**Supporting Information Figure S4c**). Residues R124, G127, W130, I143, L169, C172 and A174, identified as chemically shifted amino acids by NMR analysis (**Figure 1d,e**), displayed the lowest scores of SASA (solvent-accessible surface area, **Supporting Information Figure S4a**), which suggests that they are buried in the protein and may be indirectly affected by the binding of CLR01 to K128, given that they are in the vicinity of that binding site (**Figure 1e**). Analysis of SASA, hydrogen bond interactions, and electrostatic potential surface in simulations of JD support K128 as the preferred CLR01 binding site (**Supporting Information Figure S4a,e,f)**. Indeed, the surface charge of JD is largely asymmetric, and the region surrounding K128 is embedded in a surface patch rich in positive charges, which likely justifies the initial recruitment of CLR01 driven by favorable electrostatic interactions (**Supporting Information Figure S4f**). Once K128 is encapsulated, CLR01 initial binding is further stabilized by additional specific interactions with nearby residues. Despite being thermodynamically favored, the binding to other surface-exposed lysine residues, including K85 in the aggregation-prone region, is impeded by the formation of salt bridges and hydrogen bond interactions with neighboring residues (**Supporting Information Figure S4e**). GaMD simulations with CLR01 bound exclusively at position K128 showed a reduction of the flexibility of the hairpin compared to JD alone, as shown in the RMSD plot and B-factor representations of RMSF (**Supporting Information Figure S5a,b**). This reduction in flexibility, along with the movement of the hairpin towards the aggregation-prone region, leads to the formation of more compact structures, as evidenced by the clustering analysis (**Supporting Information Figure S5c)**, and in agreement with the ESI-MS data (**Supporting Information Figures S1c-e)**. Accordingly, a decrease in the SASA values was found for both regions, suggesting that the hairpin and aggregation-prone regions are near each other, at least during part of the simulations (**Supporting Information Figure S5d)**. A slight reduction in residue fluctuation was also observed in α5 (**Supporting Information Figure S5b**). This reduced fluctuation may enhance CLR01 binding to positively charged residues in this region, such as R103, which could explain the chemical shifts observed for this residue in the NMR experiment and the 1:2 Atx3:CLR01 complexes observed by ESI-MS (**Figure 1b**).

Taken together, our data show that CLR01 preferentially interacts with K128 at the ^124^RKLGKQW^130^ site, spatially close to the two JD termini. We propose that this interaction could disrupt early self-assembly events in the JD through allosteric regulation of the flexibility of the helical hairpin that results in the reduced exposure of amyloidogenic helix α4.[24, 25]

### CLR01 delays Atx3 self-assembly and interferes with secondary pathways in Atx3 amyloid fibril formation

Numerous studies have delved into the molecular details of Atx3 aggregation mechanism, and the influence of the flexible tail, which contains the pathological polyQ expansion on the local dynamics of the aggregation-prone JD.[23–25] Given the ability of CLR01 to interact with JD modulating the flexibility of regions critical for Atx3 self-assembly, we asked if it would affect Atx3 amyloid fibril formation. Under our experimental conditions, the Atx3 variant carrying an expanded polyQ tract (Atx3 77Q) undergoes the same amyloid self-assembly pathway as the wild-type protein (Atx3 13Q),[23, 37] as evidenced by the comparable thioflavin T (ThT) progress curves (**Figure 2a,b**). The addition of a 5-fold molar excess of CLR01 resulted in an extension of the lag phase for both Atx3 variants, a 3-fold increase in the half-time of the amyloid growth phase (t_50_), and a 4 to 5-fold decrease in aggregation rate (v_50_), as illustrated in **Figure 2** and summarized in **Table 1**. For the Atx3 13Q, the inhibitory effect of CLR01 reached a plateau at the 5-fold molar ratio, whereas a 10-fold molar ratio of CLR01 was required for Atx3 77Q to reach equivalent values for the half-time reaction coordinates (**Table 1)**. The increased lag-phase duration and decreased aggregation rates were also observed for the JD in the presence of CLR01 (**Supporting Information Figure S6a**). The control molecule CLR03, which cannot capture lysine or arginine side chains, had no major effect on Atx3 aggregation kinetics (**Figures 2a,b**, **Supporting Information Figure S6a**). The delay in the early steps of Atx3 self-assembly in the presence of CLR01 was further corroborated by the results of a Photo-Induced Cross-linking of Unmodified Proteins (PICUP) assay (**Supporting Information Figure S6b**) and by comparing the time-dependent distribution of monomer and high molecular weight (HMW) soluble Atx3 species formed in the absence and presence of CLR01 using size-exclusion chromatography (**Supporting Information Figure S6c**).

**Figure 2.**
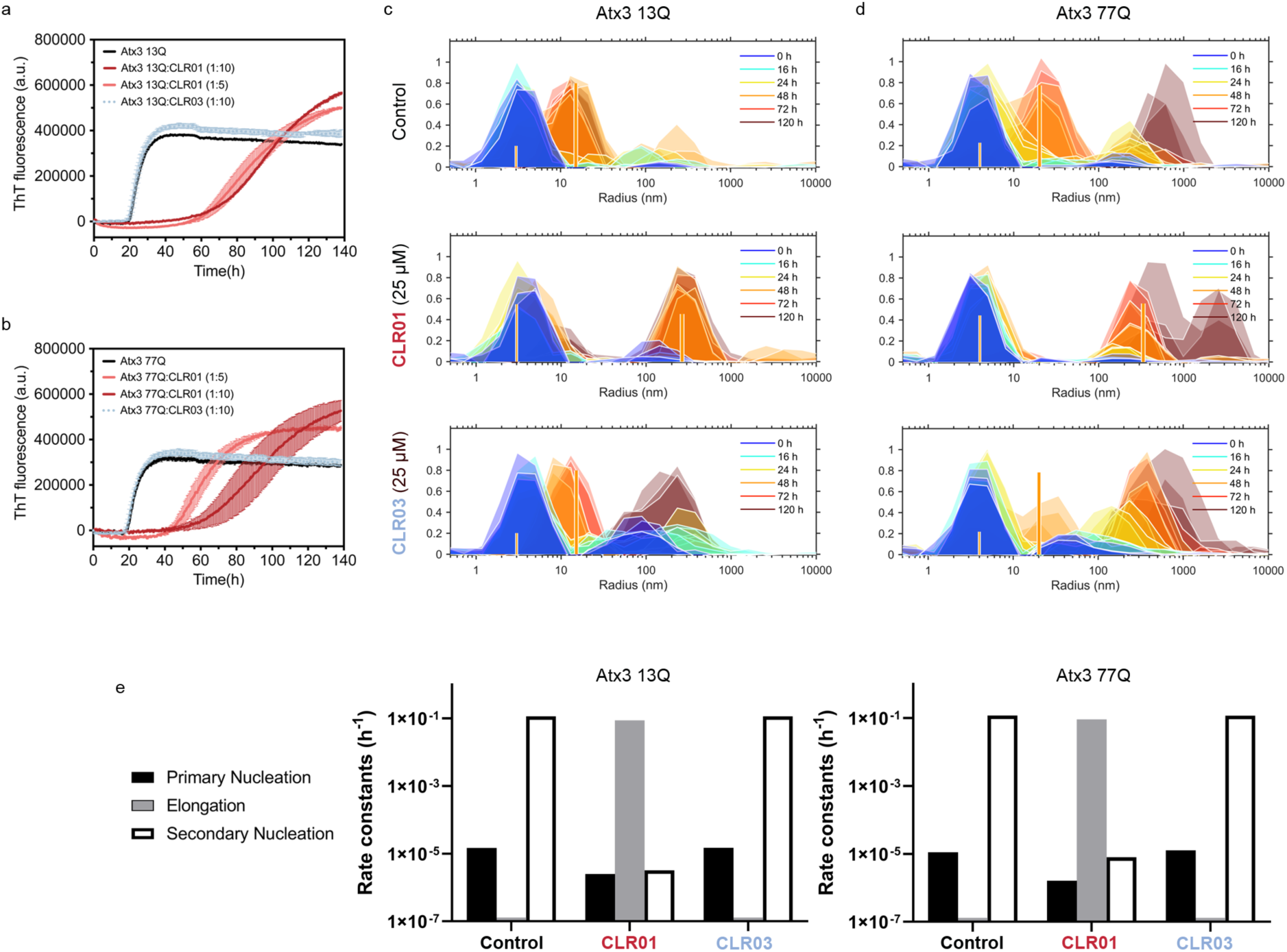
CLR01 binding modulates the Atx3 aggregation pathway, delaying amyloid fibril assembly and decreasing the secondary nucleation rate. ThT measurement of 5 μM Atx3 13Q **(a)** and Atx3 77Q **(b)** amyloid self-assembly kinetics in the absence or presence of CLR01 or CLR03. Error bars correspond to the standard deviation of three replicates. **c)** Time-dependent DLS intensity distributions during the aggregation of 5 µM wild-type Atx3 alone or in the presence of 25 µM CLR01 or CLR03. Different colors show different time points. Vertical lines: scattering intensities predicted by a nucleation-and-growth model for monomers (thinner lines) and amyloid fibrils (thicker lines) at the end of 48 h incubation (refer to **Supporting Information Figure S7** for other time points); **d)** Same as c) for Atx3 77Q. e) Fitted nucleation-and-growth rate constants (**Table 1)**.

**Table 1.**
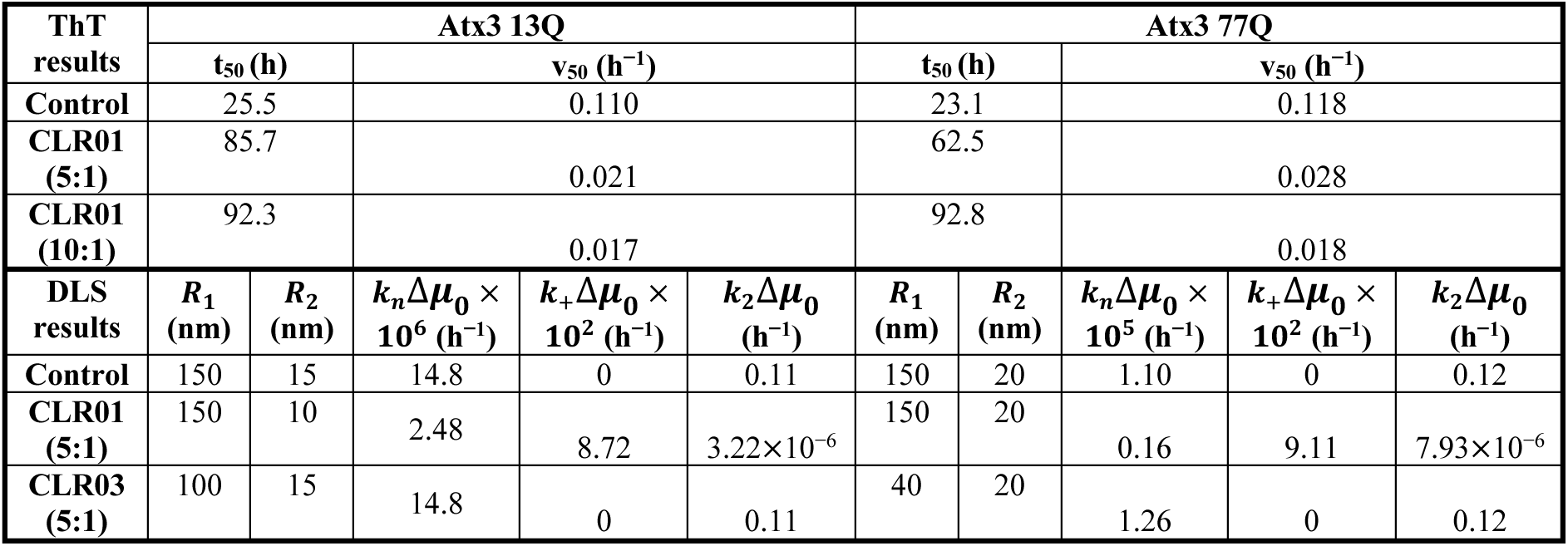
Kinetic parameters and rate constants estimated from the analysis of the ThT (top) and DLS (bottom) results.

| ThT results | Atx3 13Q |  |  |  |  | Atx3 77Q |  |  |  |  |
| --- | --- | --- | --- | --- | --- | --- | --- | --- | --- | --- |
| | $t_{50}$ (h) | $v_{50}$ ($\text{h}^{-1}$) | | | | $t_{50}$ (h) | $v_{50}$ ($\text{h}^{-1}$) | | | |
| Control | 25.5 | 0.110 |  |  |  | 23.1 | 0.118 |  |  |  |
| CLR01 (5:1) | 85.7 | 0.021 |  |  |  | 62.5 | 0.028 |  |  |  |
| CLR01 (10:1) | 92.3 | 0.017 |  |  |  | 92.8 | 0.018 |  |  |  |
| DLS results | $R_1$ (nm) | $R_2$ (nm) | $k_n\Delta\mu_0 \times 10^6$ ($\text{h}^{-1}$) | $k_+\Delta\mu_0 \times 10^2$ ($\text{h}^{-1}$) | $k_2\Delta\mu_0$ ($\text{h}^{-1}$) | $R_1$ (nm) | $R_2$ (nm) | $k_n\Delta\mu_0 \times 10^5$ ($\text{h}^{-1}$) | $k_+\Delta\mu_0 \times 10^2$ ($\text{h}^{-1}$) | $k_2\Delta\mu_0$ ($\text{h}^{-1}$) |
| Control | 150 | 15 | 14.8 | 0 | 0.11 | 150 | 20 | 1.10 | 0 | 0.12 |
| CLR01 (5:1) | 150 | 10 | 2.48 | 8.72 | $3.22 \times 10^{-6}$ | 150 | 20 | 0.16 | 9.11 | $7.93 \times 10^{-6}$ |
| CLR03 (5:1) | 100 | 15 | 14.8 | 0 | 0.11 | 40 | 20 | 1.26 | 0 | 0.12 |

Interestingly, despite the delayed aggregation kinetics, the presence of CLR01 increased the ThT fluorescence value at the end of the aggregation of JD and both Atx3 variants. The heightened ThT fluorescence signal in the presence of CLR01 is likely due to an alteration of the ThT binding pockets on the Atx3 fibril surface. Indeed, ThT is a positively charged molecule that often binds to the fibril surface via electrostatic interactions.[38, 39] Plausibly, the reversal of the positive charge of lysine/arginine by CLR01 at the fibril surface enhanced ThT binding, resulting in higher fluorescence values. It is also noteworthy that other proteins have shown an increased ThT-fluorescence signal in the presence of CLR01 even though fibril formation was decreased and CLR01 inhibited the proteins’ cytotoxicity.[40, 41]

To obtain additional mechanistic insight, the change in the size distribution of self-assembled species of Atx3 variants during the aggregation assay was analyzed using dynamic light scattering (DLS).[23, 26] First, the kinetics of the size distribution changes were measured for wild-type and Atx3 77Q (**Figure 2c, d**). The integrated analysis of both ThT fluorescence and DLS size distribution data has previously shown that the aggregation of both Atx3 variants occurs predominantly by secondary nucleation (rate constant *k_2_*), which is substantially faster than primary nucleation (rate constant *k_n_*) and fibril elongation (rate constant *k_+_*).[16, 26] The same nucleation-and-growth (NAG) model was used to describe the kinetic curves and DLS data obtained in the absence or presence of CLR01 (**Figure 2e**, **Table 1, Supporting Information Figure S7**). The fitted parameters correspond to the products *k_n_*Δ*μ*_0_, *k*_+_Δ*μ*_0_ and *k*_2_Δ*μ*_0_ (**Table 1**), where Δ*μ*_0_ is the initial supersaturation level (as detailed in **Supporting Information**). A simplified numerical fitting procedure was adopted taking into account the ThT fluorescence and DLS data. We found that the presence of CLR01 delayed protein aggregation by inhibiting the step of secondary nucleation without significantly affecting primary nucleation, which remains the rate-limiting step. Without CLR01, fibril elongation rates were too low to be quantified due to the predominance of secondary nucleation (*k_+_* << *k_2_*). For this reason, it is not clear whether the values of *k*_+_ obtained in the presence of CLR01 correspond to a modified or unmodified mechanism of fibril elongation. The rate constants *k_n_*, *k_2_* and *k_+_* that fit the DLS data also describe the half-time coordinates measured in the ThT fluorescence experiment (**Supporting Information**), further supporting the validity of our theoretical analysis.

When examining the aggregate morphologies using TEM, it became evident that CLR01 did not inhibit Atx3 aggregation but reshaped its fibrillar assembly. As depicted in **Figure 3a**, at the endpoint of the aggregation assay, when the ThT fluorescence signal reaches a plateau for both Atx3 variants (**Figure 2a,b**), Atx3 13Q formed curvilinear protofibrils with a heterogeneous distribution of lengths, whereas Atx3 77Q showed a mixture of protofibrils and fibrillar bundles corresponding to mature fibrils. In the presence of CLR01, mostly spherical oligomers (radius ∼10 nm) and small protofibrils were visible at the equivalent time points, and no mature fibrils were observed (**Figure 3a**). This is consistent with the observation of a population of very small particles with *R_H_*=10 nm in the DLS experiments for both Atx3 variants incubated with CLR01 for 72 h (**Figure 2b,c**, **Supporting Information Figure S7a,b**). Since DLS exhibits a bias toward detecting larger particles, the prevalence of a particle subset with a smaller hydrodynamic radius indicates its high abundance within the sample as clearly observed by TEM (**Supporting Information Figure 7)**.

**Figure 3.**
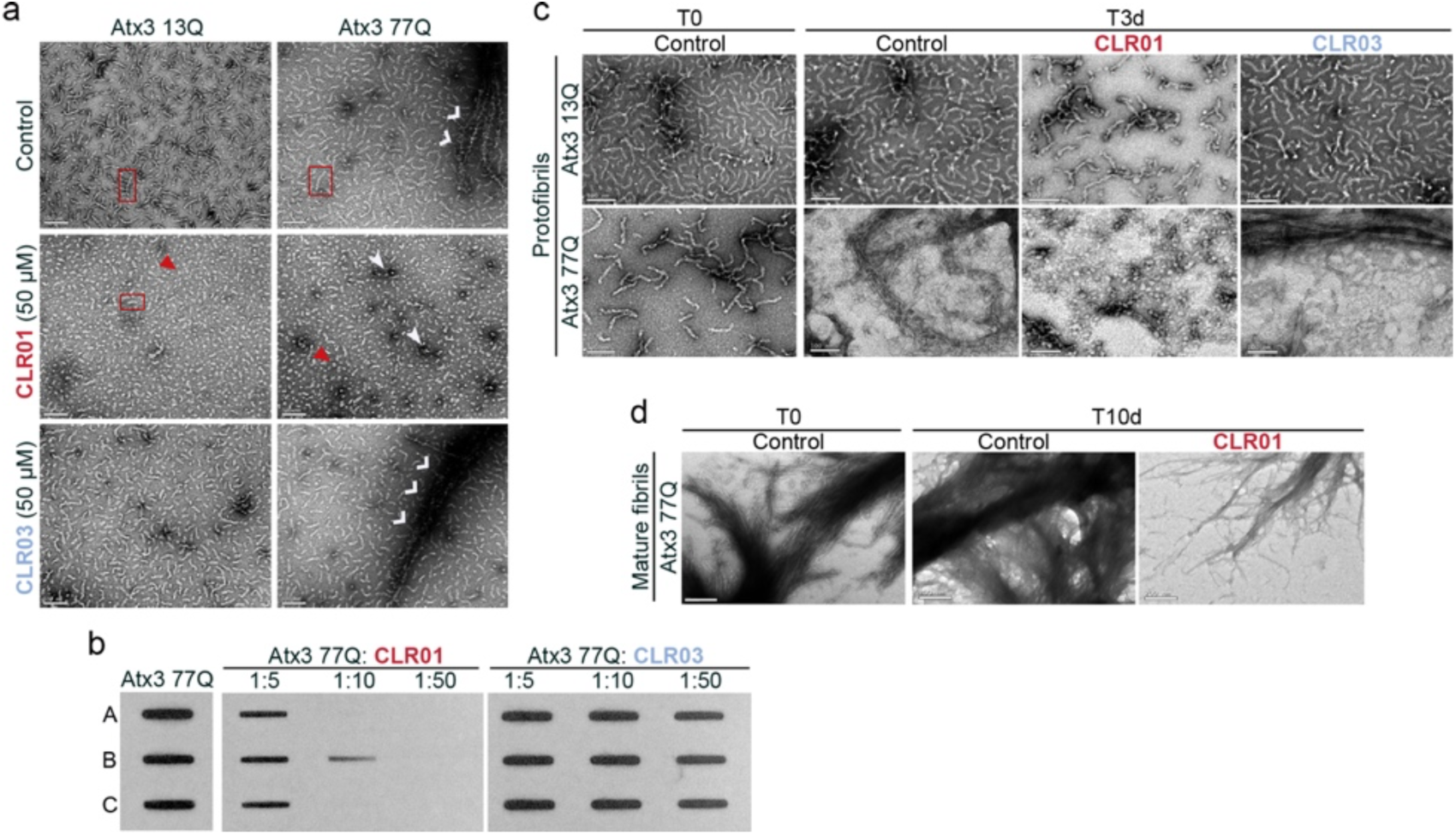
CLR01 reduces the assembly of mature fibrils of mutant Atx3 and dissociates pre-formed protofibrils and mature fibrils. **a)** Morphological analysis of 5 μM Atx3 13Q or Atx3 77Q aggregates in the absence or presence of 50 μM CLR01 or CLR03 at the endpoint of the aggregation assay, monitored by negative staining TEM. Scale bars correspond to 100 nm; orange rectangles denote Atx3 curvilinear protofibrils; red rectangles highlight small protofibrils formed in the presence of CLR01, red triangles point to Atx3 spherical oligomers; white arrowheads mark clusters of spherical aggregates; and white open arrows point to mature Atx3 Q77 fibrils. **b**) Detection of mature SDS-resistant fibrils at the endpoint of 5 µM Atx3 77Q aggregation in the presence of 5- or 10-fold molar excess of CLR01 or CLR03 by filter retardation followed by immunoblot using anti-Atx3 1H9 antibody. A, B, and C are sample replicates. **c**) Morphological analysis of 5 µM Atx3 13Q or Atx3 77Q protofibrils at t=0h and after three days of incubation at 37 °C in the absence (Control) or presence of 200 µM CLR01 or CLR0 by negative-staining TEM. Scale bars correspond to 100 nm. **d**) Morphological analysis of 5 µM 77Q mature SDS-resistant fibrils at T0h and after ten days of incubation at 37 °C in the absence (Control) and presence of 200 µM CLR01 by negative-staining TEM. Scale bars correspond to 200 nm.

The minor population of Atx3 particles with *R_H_* ≈ 150 nm detected by DLS in the presence of CLR01 is confirmed by the observation of oligomer/protofibril clusters, especially in the TEM images of Atx3 77Q species formed with 5-fold molar ratios of CLR01 (**Supporting Information Figures S7c,d**). The reduction of fibrillar bundles of Atx3 77Q by CLR01 (**Figure 3a**, **Supporting Information Figure S7c**) was supported by the decrease in mature SDS-resistant fibrils observed in a filter retardation assay (**Figure 3b**). Interestingly, high molar ratios of CLR01 were also able to disrupt pre-formed protofibrils of both Atx3 variants (**Figure 3c**) and mature fibrils of Atx3 77Q (**Figure 3d**).

Overall, these results show that CLR01 modulates the early steps of Atx3 self-assembly, including the disease-associated Atx3 77Q variant. This effect is achieved by delaying Atx3 oligomerization and the formation of protofibrils and mature fibrils through a mechanism involving a decrease in the rates of secondary nucleation. Furthermore, CLR01 can dissociate pre-formed fibril agglomerates, showcasing a multifaceted mode of action that unveils its potential for therapeutic applications in SCA3.

### CLR01 protects cortical neurons from pathological Atx3-induced synapse decline

Given CLR01’s inhibition of Atx3 self-assembly, we asked whether it could rescue the synaptic pathology in cortical neurons expressing Atx3 containing an expanded polyQ. Several studies have demonstrated that CLR01 is internalized when added to cultured neurons and is not toxic to the neurons.[42–45] First, we investigated the effect of CLR01 on the pathogenic impact of polyQ-expanded Atx3 expression in excitatory synapses of cortical neurons. Cells were transfected with Atx3 constructs containing 28Q (wild-type) or 84Q (pathological disease-associated variant) on day in-vitro (DIV) 9-10 and fixed on DIV 16. In line with previous findings,[46] the expression of the disease-associated Atx3 84Q variant led to a decrease in the levels of excitatory synapses, evident by the reduction in the intensity of the postsynaptic marker PSD95 colocalized with the presynaptic marker vGLUT1 when compared to cells expressing wild-type Atx3 (**Figure 4a,b)**.

**Figure 4.**
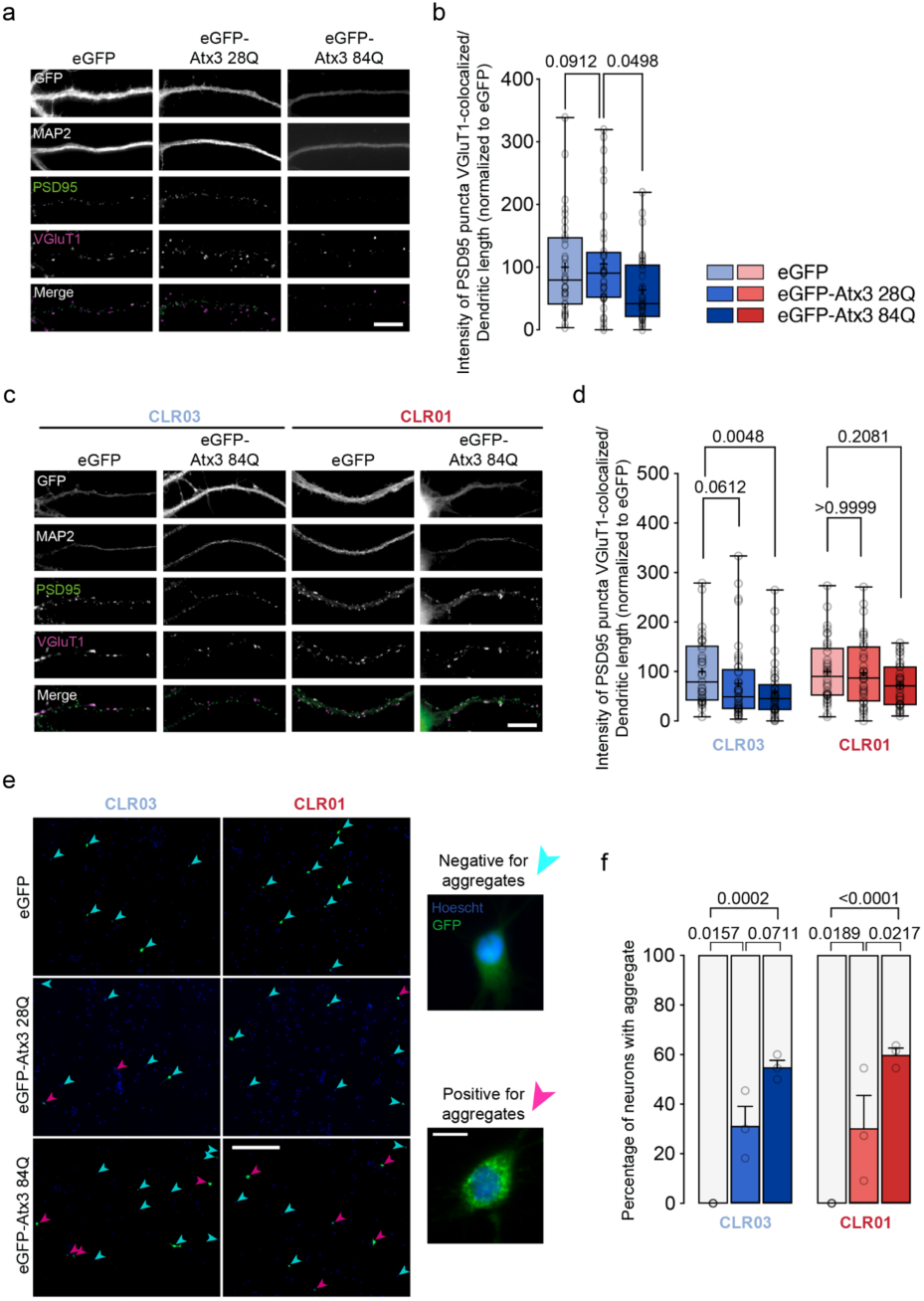
CLR01 reverts the loss of glutamatergic synapses in cortical neurons caused by pathological Atx3. Rat cortical neurons were transfected with plasmids encoding eGFP, wild-type eGFP-Atx3 28Q, or mutant eGFP-Atx3 84Q. **a**) Neurons were immunolabeled for MAP2, PSD95, and vGLUT1. Excitatory synapses were detected as PSD95 puncta that colocalized with vGLUT1; scale bar = 10 µm. **b**) Integrated fluorescence intensity of PSD95 puncta colocalized with vGLUT1 (n= 33-36 neurons per condition in 3 independent experiments). Statistical analyses were performed using Kruskal–Wallis and Dunn’s post hoc test (**Supporting Information Table S2**)**. c**) Neurons incubated with 10 µM CLR03 or CLR01 were immunolabelled for MAP2, PSD95 and VGluT1 clusters; scale bar = 10 µm**. d**) Integrated fluorescence intensity of PSD95 puncta colocalized with VGluT1 (n=35-41 neurons per condition in each of 3 independent experiments. Statistical analyses were performed using Kruskal–Wallis and Dunn’s post hoc tests (**Supporting Information Table S3**). In panels c) and d), boxes show the 25th and 75th percentiles, whiskers range from the minimum to the maximum values, and the horizontal line shows the median value. Mean is represented by “+”. **e**) GFP-Atx3 accumulates in the cell body of a fraction of transfected cortical cultures treated with CLR03 or CLR01; scale bars= 300 μm, 15 μm. **f**) Percentage of cells with GFP-Atx3 aggregates (n=3 independent experiments, 31-35 neurons per condition). Statistical analyses were performed using Two-way ANOVA and Sidak’s multiple comparisons post hoc tests (**Supporting Information Table S4**). p-values are indicated in the graphs shown in panels b), d), and f). Data are represented as mean ± SEM.

Incubation with CLR01 for 24 h rescued the PSD95/vGLUT1 colocalization levels, whereas the negative-control compound CLR03 did not (**Figure 4c,d**). Importantly, CLR01 incubation did not change the labeling for PSD95 in control neurons (**Supporting Information Figure S8a,b**). Next, we assessed whether the reversal of synaptic pathology induced by CLR01 was linked to a decrease in Atx3 aggregation within neurons. To identify Atx3 aggregates, cortical neurons transfected with eGFP-tagged Atx3 84Q were examined for eGFP-signal accumulation in the absence or presence of CLR01 or CLR03. Cells were categorized as displaying eGFP-positive aggregates if they exhibited at least one observable eGFP-positive accumulation, regardless of size. Cells showing a diffuse eGFP signal were classified as lacking aggregates. The results showed no changes in the number of cells containing Atx3 aggregates upon incubation with CLR01 (**Figure 4e,f**).

In summary, these results show that CLR01 decreased the excitatory synaptic protein content in cortical neurons expressing pathological Atx3. Interestingly this effect was not due to a reduction in the number of cells with microscopically-visible Atx3 aggregates, suggesting that these species are not toxic and do not impact synapse integrity.

### CLR01 treatment reduces neurotoxicity in a *C. elegans* model of SCA3

To evaluate if CLR01 could ameliorate disease-related phenotypes *in vivo*, we used a *C. elegans* model of SCA3, which expresses human Atx3 protein with a polyQ tract length of 130Q in the *C. elegans* nervous system and exhibits age-related locomotion defects, mimicking key features of the disease.[47] First, a food clearance assay was used to identify non-toxic concentrations of CLR01 in wild-type nematodes. None of the CLR01 concentrations tested (0.0001-100 µM) had an impact on the growth or survival of the animals after six days of exposure (**Supporting Information Figure S9**). Next, we tested the effect of chronic pre-symptomatic treatment with the same CLR01 concentrations in the *C. elegans* model of SCA3. As illustrated in **Figure 5a**, treatment with CLR01 significantly improved the animals’ motor impairments, 0.1µM being the most effective dosage (**Supporting Information, Tables S6, and S8**). Importantly, treatment with a similar concentration of CLR01 had no impact on the motility of wild-type animals, showing the specificity of the effect (**Figure 5b**).

**Figure 5.**
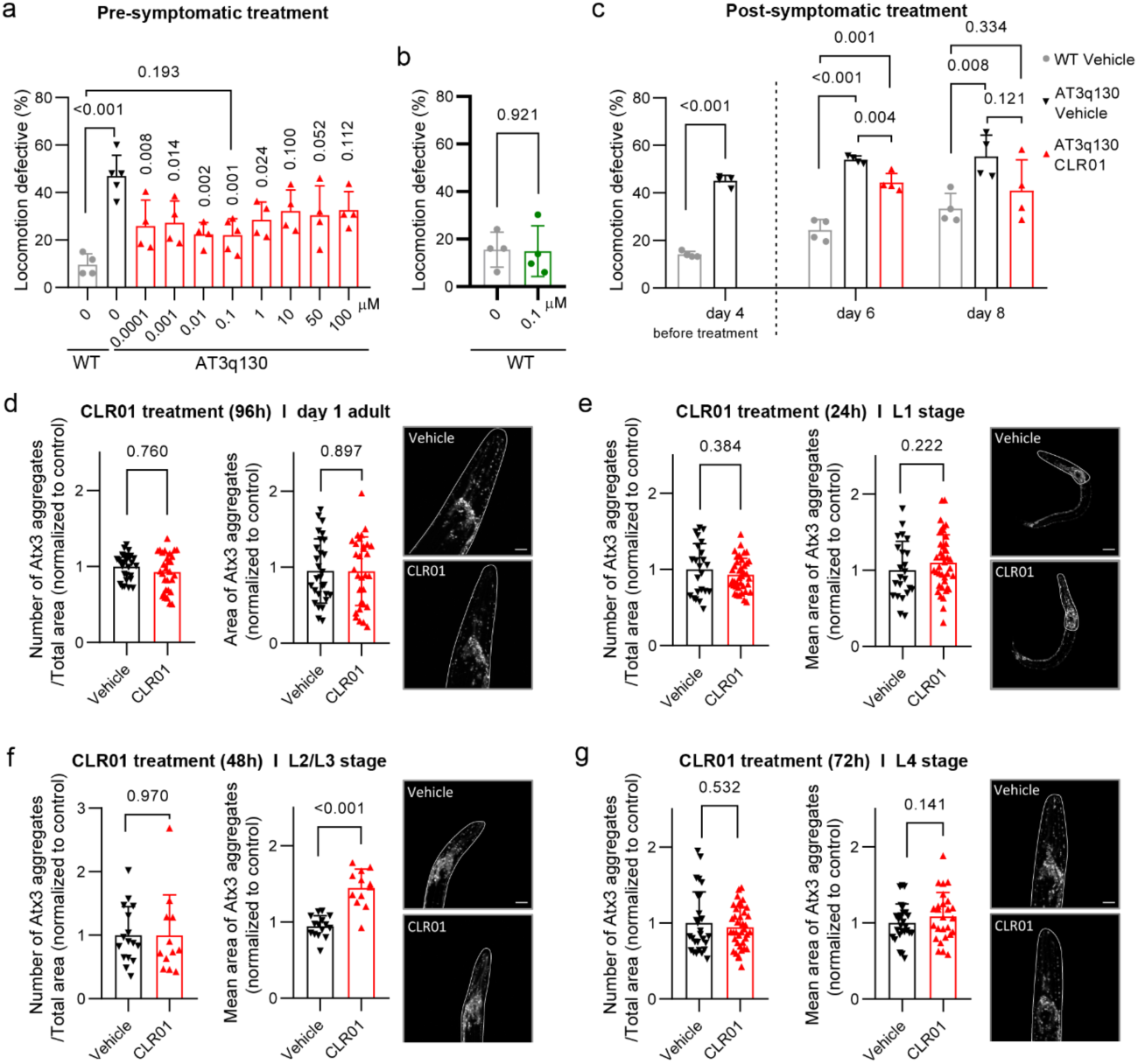
CLR01 treatment alleviates motor deficits in a *C. elegans* model of SCA3 even when treatment is initiated at a post-symptomatic disease phase. **a)** CLR01 treatment improved motor dysfunction of mutant Atx3 animals. AT3q130 animals were treated with different concentrations of CLR01 for four days, and the percentage of locomotion-defective animals was determined. CLR01 significantly impacted motor phenotype between 0.0001 and 1 µM; the maximum efficiency (%) was observed at 0.1 µM at which 67% phenotypic rescue was achieved compared to the control strain (**Supporting Information Table S6**). **b**) Treatment with 0.1 μM CLR01 for four days did not impact motor phenotype in the wild-type strain (N2). **c**) Post-symptomatic treatment with CLR01 improves the animals’ motor defects after 2 or 4 days of treatment (days 6 and 8 after hatching). (**a-c**) Data are represented as mean ± SD of 4 independent experiments, with ∼50 animals per condition/per assay (total number of animals of ∼250). A one-way ANOVA test was applied, followed by a bilateral Dunnet test for post hoc comparisons (a and c). In the comparison between the two groups (b, and c-before treatment), independent-samples t-test was used (**Supporting Information Table S7 and S8**). **d-g**) Impact of early life chronic CLR01 treatment on mutant human Atx3 aggregation pattern, labeled with an Atx3-specific antibody in the *C. elegans* model of SCA3 throughout animals’ development and adulthood. Pictures of the animal’s heads were obtained by Confocal Microscopy (Olympus FV1000) and analyzed using the MeVisLab software [48]. The graphs show a pool of 3 independent assays, with at least four animals per condition in each trial. P-values were calculated using independent-samples t-test **(Supplementary Information Table S8**). Scale bars = 20 µm.

Since the most frequent clinical situation in SCA3 is symptoms-driven genetic testing and diagnosis, we subsequently tested whether CLR01 administration would also be effective when treatment is initiated after the onset of motor impairment. Notably, even when treatment was initiated after the symptoms’ onset, CLR01 still reduced the animals’ motor dysfunction, observed by a partial reduction of the motility impairments after two days of treatment. This decreased disease progression and severity in the animals, as shown in **Figure 5c** (**Supporting Information Table S7**). The reversal of SCA3-related symptoms *in vivo* by CLR01, even upon post-symptomatic treatment, is consistent with its ability to disrupt pre-formed fibrillar bundles of pathological Atx3 (**Figure 3c, d)**. In agreement with *in vitro* ThT and TEM data (**Figures 3a-c**) and with the results obtained in cultured cortical neurons (**Figure 4e,f**), CLR01 treatment did not impede the aggregation of pathological Atx3 in SCA3 animals (**Figure 5d-g**, and **Supporting Information Table S8**) but improved disease-related motor impairment in the *C. elegans* SCA3 model, suggesting that the Atx3 aggregates formed upon treatment with CLR01 have reduced toxicity. Fluorescent aggregated foci containing Atx3 were observed within 24 h and remained unchanged by CLR01 treatment up to animal adulthood. Interestingly, the number of aggregates formed during the treatment did not significantly change after two days. However, the size of the aggregate clusters temporarily increased before decreasing again during the treatment. These results are in line with CLR01’s ability to reduce secondary nucleation rates and, in the long term, also disrupt amyloid fibrils *in vitro*, making it a promising candidate for further investigation and pre-clinical testing.

### CLR01 treatment delays disease onset and improves motor deficits in a SCA3 mouse model

Next, we asked if CLR01 treatment could ameliorate disease-related phenotypes in a mammalian SCA3 model. Taking advantage of the CMVMJD135 transgenic mouse model of SCA3 (SCA3 mice), in which quantifiable features recapitulating the motor symptoms observed in SCA3 patients can be assessed,[6] we conducted pre-clinical experiments testing the therapeutic effects of CLR01, which was previously shown to be able to cross the blood-brain barrier and be internalized by neuronal cells.[43, 45] This mouse model is well-established and has been widely used to test candidate therapies for SCA3. It presents several behavioral and neuropathological traits that closely mimic the human condition. Because CAG repeat instability is observed in the SCA3 mouse, similar to that observed in humans[49], it is important to control the number of CAG repeats within the different experimental groups. Thus, we verified that no differences existed between the mean CAG repeat number in SCA3 vehicle (VEH)- and CLR01-treated groups (**Supporting Information Figure S10a**), allowing us to avoid CAG repeat number as a confounder.[50] In CMVMJD135 mice, the onset and progression of the symptoms occur gradually during the life of the animal, the first symptoms appearing at week 6, manifesting as loss of limb strength, followed by other motor deficits that worsen with age being the motor phenotype fully established from week 16 onwards. The neuropathological findings, including mutant Atx3 nuclear inclusions and neuronal loss, occur later in life, around 22 weeks, and progress over time. In this study, CLR01 treatment was initiated early in life, at four weeks, when neurological symptoms are not yet detected in the SCA3 mice, and lasted 18 weeks (**Supporting Information Figure S10b**)[6, 8].

Chronic treatment with CLR01 caused no detrimental effects to WT mice, as their body weight (**Supporting Information Figure S10c**), fur quality, activity, presence of grooming, and nesting behaviors were comparable to those of wild-type mice receiving vehicle. CLR01 treatment delayed the reduction of body weight seen in the SCA3 mice, a two-week delay being observed; nevertheless, at 12 weeks of age the body weight loss reached the SCA3 vehicle group level, suggesting that this effect on body weight was not sustained throughout time (**Supporting Information Figure S10c**). No evident toxicity was observed, as CLR01-treated mice performed similarly to the vehicle-treated mice in all the motor tests, suggesting a safe profile for CLR01 chronic treatment in these mice, similar to previous models.[28] CLR01 treatment showed a strong therapeutic effect on early motor symptoms in the SCA3 mice, including a delay in the symptom onset and overall improved performance in the swimming test (**Figure 6a**) and to a lesser extent in the balance-beam test, in which at 12 weeks of age the CLR01-treated SCA3 mice performed indistinguishably from WT mice, but by 16 weeks of age, their performance deteriorated and was similar to that of vehicle-treated SCA3 mice (**Figure 6b**). Importantly, CLR01 administration substantially improved the gait quality of SCA3 mice and delayed the appearance of an abnormal gait pattern by eight weeks (**Figure 6c**).

**Figure 6.**
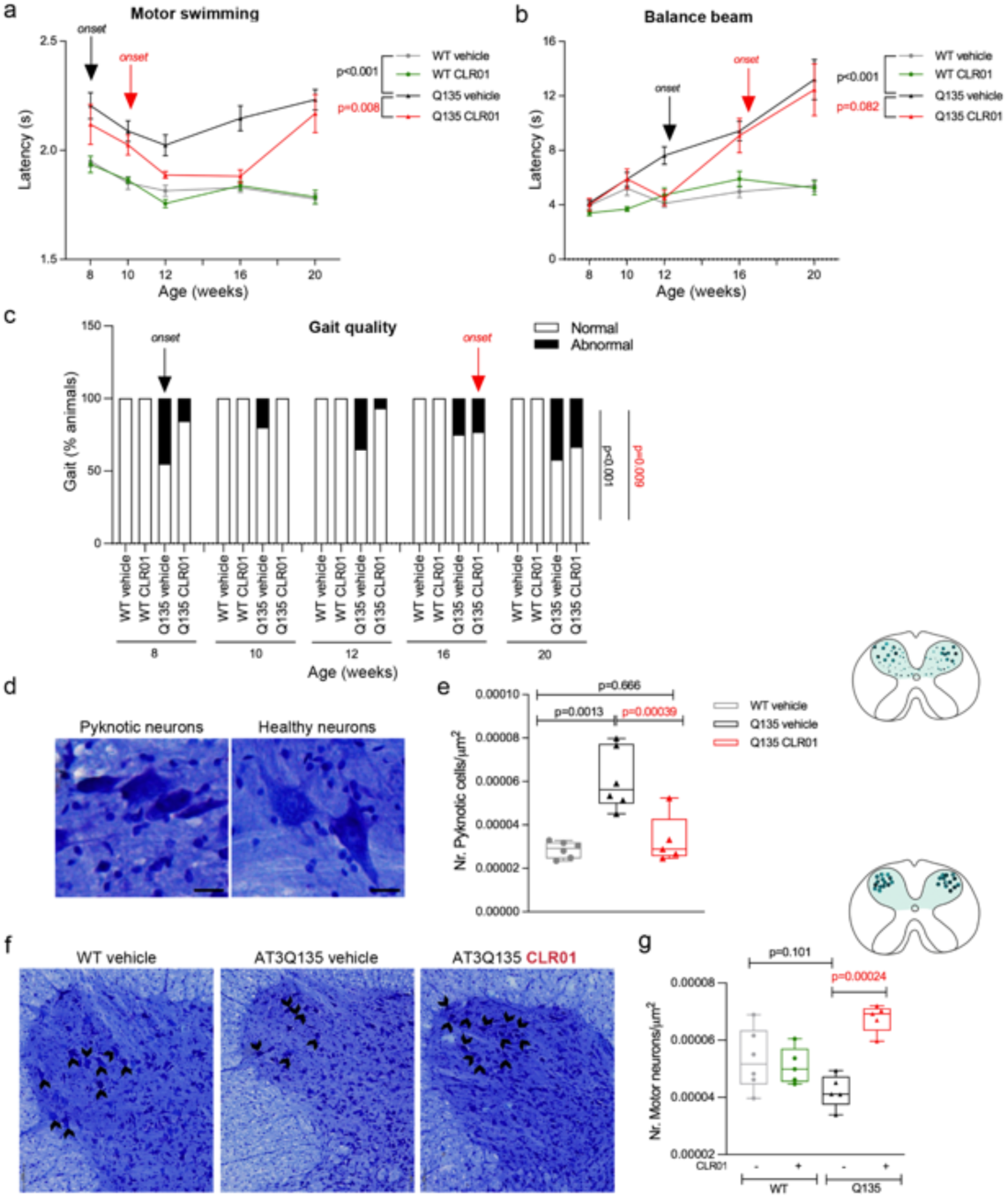
CLR01 chronic administration improves motor function and neuropathology of SCA3 mice. **a**) the latency to cross a swimming tank, representing both the strength and the movement coordination ability of the animals, is higher in vehicle-treated SCA3 mice. The severity of this symptom improves over time with CLR01 treatment, which has a delayed onset and is effective up to 16 weeks of age.; **b**) CLR01 improves SCA3 mice balance, decreasing their latency to cross a 12 mm-squared beam. **c**) The gait quality of SCA3 animals is strikingly improved by CLR01 treatment, delaying the onset of ataxic gait by eight weeks. **d**) Representative images of Cresyl Violet staining in the cervical spinal cord of healthy and pyknotic cells. **e**) Vehicle-treated SCA3 mice showed a substantial increase in the number of pyknotic cells, which was rescued by daily CLR01 administration for 18 weeks. **f**) Representative pictures of the motor-neuron cell bodies (black arrows) in the ventral horn of the cervical spinal cord. **g**) CLR01 treatment increases the number of motor neurons in the cervical spinal cord. Sample size and statistical details are provided in **Supporting Information, Table S9**. The scale bar for lower magnification pictures represents 200 μm, and for higher magnification, 100 μm. P-values are shown in the graph. Black lettering represents the comparisons between WT-vehicle and SCA3-vehicle. Red lettering represents the comparisons between SCA3-vehicle and SCA3-CLR01.

Next, we asked if the neuropathology features of SCA3 mice were improved by CLR01 treatment. We have shown previously that pyknosis, a process in which the cell nucleus becomes dense and compact, indicating the initiation of a degeneration process, occurs in the brain of the SCA3 mice.[6, 8] CLR01 chronic treatment fully rescued this degenerative process in the cervical spinal cord of SCA3 mice by reducing the number of pyknotic cells to vehicle-treated WT mice levels (**Figure 6d, e**). Previous studies have shown that at 34 weeks of age cholinergic motor neurons degenerate in the spinal cord of SCA3 mice.[6, 8] Interestingly, CLR01 treatment led to the preservation of cervical spinal cord neurons in SCA3 mice and did not impact the number of these neurons in WT animals (**Figure 6f, g**). Additionally, CLR01 administration had no major impact on the number of Atx3-positive nuclear inclusions (**Supporting Information Figure S11**), which are usually found in the brain and spinal cord of SCA3 mice. [6, 8, 51] As the presence of these end-stage aggregates in CLR01-treated mice is associated with improved histopathological findings, we propose that the detected Atx3 aggregated species have a reduced proteotoxicity.

## DISCUSSION

Protein misfolding and its subsequent assembly into amyloid and amyloid-like deposits represent distinctive features consistently observed in numerous severe neurodegenerative conditions, such as Alzheimer’s, Parkinson’s, and Huntington’s diseases. In SCA3, similar to other polyglutamine expansion disorders, intranuclear inclusions immunopositive for Atx3 are found in neurons.[4, 52, 53] Although the exact role of Atx3 self-assembled molecules in causing selective neuronal demise has not yet been fully deciphered,[54, 55] several approaches to reduce the aggregate burden have been pursued as strategies for the development of disease-modifying therapies for SCA3. Here, we show that CLR01, a supramolecular ligand with an affinity for lysine and, to a lesser extent, arginine residues,[28, 56] modulates the aggregation behavior of Atx3 *in vitro* and has a positive impact in reverting disease-related phenotypes in SCA3 cell and animal models though it does not reduce the neuronal aggregates. This suggests that these microscopically-visible aggregates formed in the presence of CLR01 are unlikely to be toxic. It remains to be determined whether CLR01 promotes the formation of non-toxic Atx3 aggregate polymorphs with structural characteristics that differ from those formed in the absence of the molecular tweezer.

### An optimal chemical environment drives CLR01 recruitment to the Josephin domain

In silico and in vitro studies showed that CLR01 selectively binds to the globular JD, a key player in initiating the early stages of fibrillar aggregate formation. To explore the interaction mechanism, we combined NMR, which is highly sensitive to changes in the local chemical environment of the ligand binding sites, with extensive enhanced sampling MD simulations that can capture transient interactions while sampling various conformational states. We showed that CLR01 forms a complex with K128, located in a positively charged region in the vicinity of an exposed hydrophobic surface patch (**Figure 7**). This region displays an ideal chemical environment for the establishment of favorable interactions with CLR01: while encapsulating K128, the phosphate groups of the molecular tweezer can engage into electrostatic interactions, while its convex surface may be further stabilized through hydrophobic interactions with neighboring hydrophobic residues. Upon binding to K128, CLR01 modulates the conformational fluctuations in the distal helical hairpin, through allosteric long-range effects that restrict the local misfolding necessary for exposing the amyloidogenic region in helix α4, which initiates Atx3 self-assembly.

**Figure 7.**
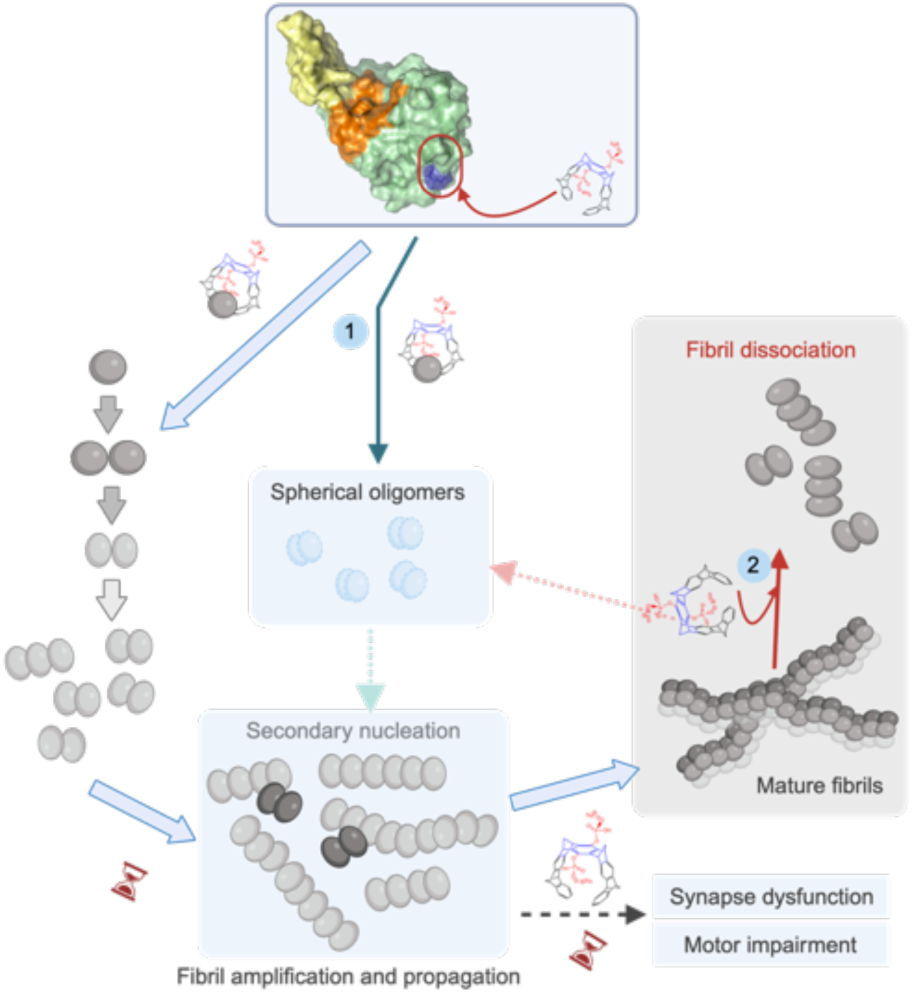
Schematic representation of the dual action of CLR01 on the delay of Atx3 self-association (1) and fibril disruption (2). Binding of CLR01 to K128 (colored dark blue) within a positively charged surface patch opposite to the aggregation-prone region (orange) triggers a conformational change in the helical hairpin (yellow) of the Josephin domain. The CLR01-induced conformational change reshapes the Atx3 aggregation pathway and reduces secondary nucleation rates. This modulation of the self-assembly pathway correlates with the reversion of synapse dysfunction in SCA3 cell models and a delay in the onset of motor impairment in SCA3 animal models. The spherical oligomers observed upon incubation with CLR01 may be on-pathway to protofibril assembly or result from fibril dissociation processes. Created in BioRender (https://BioRender.com/e62f696).

### CLR01 decreases secondary nucleation processes, modifies Atx3 fibril assembly and dissociates pre-formed amyloid fibrils

The analysis of Atx3 aggregation in the presence of CLR01 revealed a pronounced kinetic impact on amyloid fibril assembly. Our in vitro assays confirmed that CLR01 effectively interferes with the assembly of Atx3 into amyloid structures and delays the formation of mature fibrils of the pathological Atx3 77Q. We have shown previously that surface-mediated secondary nucleation prevails in the in vitro kinetics of Atx3 fibril formation of both wild-type and disease-related variants.[16, 26, 27] CLR01 specifically decreases the rate of secondary nucleation and thereby restricts the autocatalytic amplification of Atx3 amyloid species. Several scenarios can be put forward to explain the impact of CLR01 on these secondary nucleation processes. One possibility is that CLR01 interaction may restrict the local misfolding and the ability of Atx3 monomers to adopt the structure required for nucleation and assembly at the surface of pre-formed fibrils. Alternatively, it may directly impede the binding of monomers/oligomers to the fibrils or modify the properties of the parent fibrils required for the attachment step. Notably, CLR01 does not completely prevent Atx3 fibril formation, and mature fibrils of the disease-related variant eventually appear. Intriguingly, CLR01 incubation gives rise to a distinct population of smaller oligomers compared to the Atx3 fibrils formed in the absence of the molecular tweezer. Whether these are on-pathway amyloid species or structurally distinct Atx3 fibril polymorphs that may later convert to extended fibrillar aggregates with higher thermodynamic stability[57] remains to be elucidated. Future studies addressing a time-resolved structural analysis of fibril polymorphs formed along the aggregation pathway, similar to the recent structural analysis of tau[58] and human islet amyloid polypeptide[57] will help to address these questions. It is also noteworthy that CLR01 dissociated pre-formed Atx3 protofibrils and mature fibrils, exhibiting a fibril-disrupting activity that has also been shown for α-synuclein[59] and Aβ peptide[44] aggregates. This is a slow process that requires a 40-fold molar excess of the molecular tweezer, but it suggested that CLR01 could be used as a treatment not only in the early but also at later stages of the disease. The exact mechanisms that lead to the disruption of the Atx3 fibrils have yet to be fully understood. However, we speculate that at relatively high molar concentrations, CLR01 may interact with weaker affinity binding sites containing exposed positively charged residues in the C-terminal tail of Atx3 and weaken the intermolecular interactions that stabilize the fibrillar assemblies.

### CLR01 reverts SCA3-related phenotypes in cell and animal disease models

The thorough mechanistic analysis of CLR01’s impact on Atx3 aggregation, particularly its role in reducing secondary nucleation, a process closely associated with neurotoxicity and the proliferation and spreading of amyloid fibrils[60, 61], prompted the investigation into its effects in SCA3 cell and animal models. One key question is how the mechanisms for aggregation impairment dissected in vitro translate into tangible improvements of the pathology in vivo. Importantly, in our SCA3 neuronal model, CLR01 treatment rescued the synapse pathology occurring when cultured cortical neurons express pathological Atx3 with an expanded polyQ tract. Importantly, CLR01 restored motor dysfunction in the SCA3 *C. elegans* model and showed a clear therapeutic effect in SCA3 mice. In this context, CLR01 delayed the onset of motor symptoms and rescued spinal cord and motor-neuron degeneration. It is noteworthy that the efficacy of CLR01 in reversing or delaying SCA3-related phenotypes in cultured cortical neurons and *in vivo* was not accompanied by a reduction in microscopically visible Atx3 aggregates. In fact, the identification of Atx3 deposits in both affected and non-affected brain regions of patients who died of SCA3 has suggested that these microscopically-visible inclusions may not represent the neurotoxic entities[54, 55]. As in other amyloid diseases, growing evidence supports the idea that oligomeric and/or protofibrillar species play a critical role in triggering and spreading polyQ pathology in neurons[62–64]. The beneficial effects observed combined with the presence of large aggregates in SCA3 cells and animal models suggest that in cultured neurons and in vivo, CLR01 may indeed interfere with secondary nucleation processes responsible for the autocatalytic proliferation and spreading of toxic Atx3 aggregates, which are stealth to the applied imaging methodologies. In support of this hypothesis, treating SCA3 mice with CLR01 from an early disease state protected the neuronal dysfunction, enabling their preservation even after symptom installation.

### The temporary efficacy of CLR01 treatment in the SCA3 mouse model

An intriguing observation in the mouse model is that despite a delayed onset of several disease-related motor symptoms, once the symptoms appear, they evolve rather quickly to levels that mirror those of the untreated SCA3 mice. Several hypotheses may be put forward to explain this observation. First, one may argue that CLR01 reshapes the Atx3 aggregation pathways, leading first to the appearance of non-toxic Atx3 amyloid intermediates, which are kinetically favored. Later in the aggregation process, these may convert to thermodynamically more stable fibril polymorphs that are competent to template the assembly of new fibrils, which grow exponentially, accumulate, and spread in the mouse brain and spinal cord, resulting in motor dysfunction. Alternatively, the continuous subcutaneous administration of CLR01 could result in the accumulation of the molecular tweezer in the mouse brain.[42] This build-up may drive CLR01 interactions with weaker affinity binding sites on Atx3, including lysine residues necessary for Atx3 post-translational modifications[65–68] and nucleocytoplasmic shuttling,[69, 70] with implications on protein stability aggregation and turnover.[71–73] The interaction of CLR01 with lysines/arginines on secondary binding sites might also justify the decreased effect of CLR01 treatment on locomotion defects in *C. elegans* observed at concentrations higher than 1 µM. The transient beneficial effects of CLR01 treatment in the mouse model suggest that the molecular-level mechanistic information gathered in this study will be critical for developing improved lead molecules for effective SCA3 therapies.

### CLR01 as a scaffold for allosteric modulators of pathological Atx3 aggregation

In conclusion, SCA3 is a severe neurodegenerative disease with an unmet and urgent need for disease-modifying therapies. Numerous studies have demonstrated the efficacy of therapeutic strategies such as silencing the pathological polyQ-expanded Atx3 or addressing downstream proteotoxicity.[74, 75] We have recently shown that increasing dopamine levels offers a promising approach to stopping disease progression by interfering with the secondary nucleation step of Atx3 aggregation.[16] However, the lack of a comprehensive analysis of their mechanisms of action has hindered the translation of these findings into clinical applications. This study presents a systematic multiscale analysis of CLR01 efficacy, spanning from the molecular level to the SCA3 mouse model. Our results demonstrate that CLR01 targets the globular domain of Atx3, limiting the exposure of the aggregation-prone site, which is crucial for initiating early aggregation steps. Furthermore, CLR01 interferes with the autocatalytic proliferation of Atx3 fibrils through secondary nucleation, an effect that is translated into the overall improvement of SCA3 phenotypes in cultured neurons and in animal models. Notably, CLR01 delays the onset of motor function loss in a pre-clinical SCA3 mouse model, hinting at its potential as a disease-modifying therapy for SCA3 patients, which could be used in combination with other therapeutic approaches currently under development. Our results offer crucial mechanistic insights that advance our comprehension of the self-assembly pathways of Atx3 in SCA3 and underscore the role of the distal termini of the JD in shaping its conformational landscape and aggregation propensity. A previously undisclosed allosteric site distant from the ubiquitin substrate binding region was identified, providing critical knowledge for designing tailored molecular tweezers capable of precisely targeting the pathological aggregation routes while preserving the normal function of the native protein in cell proteostasis. These findings expose novel opportunities to identify allosteric binding pockets in “undruggable” aggregation-prone proteins associated with multiple human diseases, whose self-assembly regions are highly flexible and challenging to target.

## EXPERIMENTAL SECTION

Atx3 variants and compounds. Human full-length Atx3 (Uniprot P54252-2) containing 13 or 77 glutamines within the polyglutamine tract (Atx3 13Q, MW of 43582.5 Da or Atx3 77Q, MW of 51882 Da, respectively), and the truncated variant comprising the N-terminal globular Josephin domain (Atx3 JD, MW of 23767.87 Da) were used in the experiments, all containing a N-terminal 6xHis-tag. The details about the expression vectors used are in the plasmid repository database Addgene with the accession codes 184247, 184248, and 184249. CLR01[29] and CLR03[76] were provided by Gal Bitan’s lab.

### Protein purification

Atx3 variants were expressed in *E. coli* and purified by metal-affinity chromatography and size-exclusion chromatography as described previously.[23, 26, 37] The protein concentration was determined by measuring the absorbance at 280 nm using the extinction coefficient of 35660 M^-1^.cm^-1^ (Atx3 JD) or 31650 M^-1^.cm^-1^ (Atx3 13Q and Atx3 77Q). The purified proteins were stored at -80 °C in 20 mM sodium phosphate pH 7.5, 150 mM NaCl, 5% (v/v) glycerol, 2 mM EDTA, 1 mM DTT.

### NMR titrations

SOFAST-HMQC NMR spectra were recorded at 21 °C on a 800-MHz Bruker Avance instrument (Bruker, Germany) equipped with a cryogenic probe. Samples of ^15^N-labelled JD were used at 100 μM after overnight dialysis in a buffer containing 20 mM sodium phosphate pH 7.5, 100 nM NaCl, and 1 mM DTT. 10% D_2_O was added before measurements. CLR01, dissolved in a matching buffer, was titrated into the JD solution to reach a final protein:CLR01 molar ratio of 1:2 in 0.25 increments. The results were analysed by comparing the chemical shift perturbation effects observed by titration of CLR01 with the spectral assignment of the isolated JD deposited in the BMRB database (entry number 6241).

### Enhanced Sampling simulations

The coordinates of the open (PDB code 1YZB[33]) and closed (PDB code 2AGA[34]) conformations of the JD were used as the initial coordinates for the GaMD simulations[30] in the absence or the presence of CLR01, as described in Supplementary Information. The CHARMM36m force field[77] was used for the protein in all simulations. The CHARMM-GUI server[78, 79] was employed for the preparation of all systems. During the Gaussian accelerated molecular dynamics (GaMD) protocol, 2.4 ns of classical MD were carried out, followed by 8-12 ns of GaMD equilibration until convergence of k0D and k0P to 1. Then, dual boost GaMD production runs were carried out (3 x 200 ns replicas) for all the setups using AMBER 20.[80] In addition, JD (open conformation) was simulated in complex with a single CLR01 molecule positioned at K128, with three replicas of 500 ns each. The CL-FEP approach[36] was employed to estimate the Gibbs energy of binding of CLR01 to K/R residues of the JD, as described in the **Supporting Information**.

### Native Mass Spectrometry

Protein solutions were buffer-exchanged into 20 mM ammonium acetate using 10 kDa Amicon filters and diluted to 10 μM. CLR01 was added to the solution at 10 μM to induce binding. Solutions containing protein and protein/CLR01 complexes were sprayed via nanospray capillaries (ThermoFisher Scientific) on a Synapt G2-Si (Waters). The instrument was run in mobility TOF mode with a capillary voltage of 1-2 kV. The sampling cone was set to 20 V and the source offset was set to 20 V. Mobility was measured over the entire m/z range and mobiligrams were extracted from the [M+nH]^n+^ peak. Mass spectra were deconvoluted using UniDec.[81]

### Atx3 aggregation assays

#### Thioflavin-T (ThT) binding assay

Immediately before each aggregation assay, purified protein solutions were applied to a Superose 12^®^ 10/300 GL column (GE Healthcare Life Sciences) pre-equilibrated in aggregation buffer (20 mM HEPES pH 7.5, 150 mM NaCl, 1 mM DTT) and 0.4 mL fractions were collected. The peak corresponding to monomeric protein was collected for further aggregation assays. ThT assays were carried out in the absence or presence of CLR01 or CLR03 at 37 °C, as described previously, in low protein-binding Corning^®^ Thermowell™ 96-well polycarbonate PCR microplates. A protein concentration of 5 µM and ThT concentration of 30 µM were used with three technical replicates.[23, 26, 37] The fluorescence (emission 480 nM, excitation 440 nm) was monitored every 30 min on a FluoDia T70 microplate fluorimeter (Photon Technology International, Edison, NJ, USA). Data were analyzed using Prism 5 (GraphPad Software). Sample quality control was determined at several time points during the aggregation assay by Transmission Electron Microscopy (TEM) using a TEM JEM-1400 (JEOL, Tokyo, Japan) at an accelerating voltage of 80 kV, as described previously. [23, 26, 37] At the endpoint of the aggregation assays, samples were analysed by a filter-retardation assay following a protocol described previously.[16, 26]

#### Atx3 oligomerization time-course analysis

The aggregation of the proteins in the absence or in the presence of CLR01 or CLR03 was evaluated by size exclusion chromatography (SEC), TEM, and dynamic light scattering (DLS). The proteins (5 μM final concentration) were incubated in aggregation buffer at 37 °C without shaking. At different time points, 50 μL aliquots were collected and analyzed directly by TEM or filtered and injected onto a Superdex 200 Increase 5/150 GL column (GE Healthcare Life Sciences) as described previously for Atx3 13Q.[16, 26] Measurements of the soluble species present in each sample (monomers, intermediate oligomeric species, high molecular weight (HMW) oligomers) was made by determining the relative peak areas (RPA) at each time point after integration of SEC chromatograms peak areas and normalization by the total area of the peaks. Measurements of DLS autocorrelation functions were carried out on a Zetasizer Nano ZS DLS instrument (Malvern Instruments) during the aggregation of 5 mM protein in the absence or presence of 5- or 10-fold molar excess of CLR01 or CLR03, as described previously[16]. The samples were incubated in a UV-Cuvette micro cuvette (Brand, Germany) with a cap, maintained at a constant temperature of 37 °C without shaking, and analyzed periodically at 0, 16, 24, 48, 72 and 120 h at 25 °C and a scattering angle of 173° to the incident beam. Intensity size distributions were obtained by CONTIN analysis of the measured autocorrelation functions using the MATLAB code rilt.[82, 83]

#### Atx3 fibril dissociation assays

At the endpoint of the aggregation assay and after quality control of fibrils by TEM (time 0 h), Atx3 13Q and 77Q protofibrils, and Atx3 77Q mature fibrils (5 μM final concentration), were incubated in the absence or presence of 40-fold molar excess of CLR01 or CLR03 in aggregation buffer at 37 °C without shaking. Fibril dissociation profile was evaluated by TEM as described above, after 0, 1, 2 or 10 days of incubation.

### Neuronal cell model

#### Primary cortical neuronal cultures

Dissociated neuronal cultures were prepared as described previously.[84] Neurons were plated in neuronal plating medium (MEM supplemented with 10% (v/v) horse serum, 0.6% (w/v) glucose and 1 mM pyruvic acid) onto poly-D-lysin-coated coverslips. Low-density cultures were plated in culture dishes at a final density of 11.6k or 13.9k cells/cm^2^. After 2-4 h of incubation at 37 °C in a humidified incubator containing an atmosphere of 5% CO_2_/95% air, coverslips were flipped over an astroglial feeder layer (Banker cultures) in NeuroBasal Medium (NBM; supplemented with SM1 (1:50 dilution), 0.5 mM glutamine, 0.12 mg/mL gentamycin). The cultures were maintained at 37 °C in a humidified incubator under 5% CO_2_/95% air, and fed once per week with fresh NBM without glutamate. To prevent glia overgrowth, neuronal cultures were treated with 10 µM FDU (5-deoxyfluoruridine) after 3 day in-vitro (DIV). All animal procedures were reviewed and approved by ORBEA and Direção Geral de Veterinária (DGAV, Portugal).

#### Transfection protocol

DNA constructs were transiently expressed in primary neuronal cultures using a calcium phosphate transfection protocol adapted from Jiang and collaborators.[85] Cultures were transfected at 10 DIV, with 3 µg of plasmids encoding eGFP, non-expanded eGFP-Atx3 28Q or expanded GFP-Atx3 84Q. All incubations were performed at 37 °C in a humidified incubator under 5% CO_2_/95% air. Plasmids were allowed to express for 6 days.

#### Immunofluorescence

Low density cultured neurons (DIV 16) were fixed with 4% paraformaldehyde (PFA) / 4% sucrose in Phosphate Buffer Saline (PBS; in mM 137 NaCl, 2.7 KCl, 1.8 KH_2_PO_4_, 10 Na_2_HPO_4_·2H_2_O, pH7.4) for 15-min at room temperature, followed by 6 sequential washes with PBS. After permeabilization with 0.25% (w/v) Triton X-100 in PBS for 5 min at 4 °C, neurons were washed with PBS and blocked with 10% (w/v) bovine serum albumin (BSA) for 30 min at 37 °C. Subsequently, cells were incubated with primary antibodies diluted in 3% BSA overnight at 4 °C or for 2 h at room temperature. To stain for excitatory synapses, anti-PSD95 antibody (mouse, 1:500, Cell Signalling Technology) and anti-VGLUT1 antibody (guinea pig, 1:1000, Merck Millipore) were used; dendrites were identified with anti-MAP2 antibody (chicken, 1:5000, Abcam). Neurons then were washed 6 times with PBS and re-incubated for 1 h with secondary antibodies in 3% BSA at 37°C. The following secondary antibodies from Invitrogen Molecular Probes were used:Alexa Fluor® 488 conjugated anti-rabbit (1:500), Alexa Fluor® 568 conjugated anti-mouse (1:500), Alexa Fluor® 647 conjugated anti-guinea pig (1:500), and AMCA-conjugated anti-chicken (1:200, Jackson ImmunoResearch). Following six washes, coverslips were mounted with fluorescence mounting media.

#### Quantification of aggregate-containing neurons

Transfected cortical neurons at DIV 16 were fixed as described for immunofluorescence. 10 to 13 neurons *per* condition were visualized using a Carl Zeiss Axio Observer Z1 microscope using a 40× oil objective (EC Plan-Neofluar, 1.3 NA), and eGFP-Atx3 aggregate-containing neurons were counted versus the number of cells presenting only diffuse eGFP-Atx3 signal, as described previously.[46] Aggregation was expressed as the percentage of cells containing aggregates.

#### Image quantification

Images were quantified using the image analysis software FIJI. Background intensity was subtracted from the user-defined threshold and normalized to the cell body area. Synapses were evaluated using a semi-automatic macro designed for this purpose. The area of interest was randomly selected by using MAP2 staining for non-transfected cells and GFP staining for transfected cells. Dendritic length was measured in the area of interest, using one of the mentioned channels. To quantify the proteins of interest, images were subjected to a user-defined intensity threshold to have defined protein clusters and user-defined background intensity was subtracted in all images. Each protein cluster present in the area of interest was analyzed and intensity, area, and number measurements were obtained. Synaptic clusters were selected by overlapping PSD-95 clusters of interest with thresholded VGluT1 clusters. All results were normalized to dendritic length. Image analysis was performed blinded to the experimental condition.

#### Treatments

Cells (DIV 15) were treated with 10 µM CLR01 or CLR03 for 24 h before fixation.

### Caenorhabditis elegans SCA3 model

#### *C. elegans* strains, maintenance and synchronization

The wild-type Bristol N2 strain was obtained from *Caenorhabditis Genetics Center* (Univ. of Minnesota). The *C. elegans* model of SCA3, expressing full-length human ATXN3 proteins with 130 glutamines [AT3q130: AM685 *rmls263(P_F25B3.3_::AT3v1-1q130::yfp)*] throughout the animals’ nervous system was generated as described previously.[47] Animals were grown at 20 °C on nematode growth medium (NGM) agar plates seeded with *Escherichia coli* (*E. coli*) OP50 strain, which was grown overnight at 37 °C and 180 rpm in Luria Broth (LB) media.[86] To obtain an age-synchronized population of eggs, gravid adults were treated with an alkaline hypochlorite solution (0.5M NaOH, ∼2.6% NaClO) for 5 minutes. The eggs were washed in M9 buffer, resuspended in S-medium to the appropriate egg number, and transferred into the 96-well plates (drug assays). Batches of OP50 bacteria were grown overnight at 37 °C and 180 rpm in LB, pelleted by centrifugation, inactivated by three cycles of freeze/thawing, frozen at -80 °C and then resuspended in S-medium supplemented with cholesterol, streptomycin/penicillin and nystatin (Sigma).

#### CLR01 preparation for *C. elegans* assays

A 50 mM stock solution of CLR01 was prepared in DMSO. For each concentration to be tested, a stock solution was prepared at a 100x concentration in 100% (v/v) DMSO (D5879-Sigma), from which work dilutions were prepared 2.4x concentrated. A final concentration of 1% (v/v) DMSO was used in all the conditions tested to avoid solvent-specific developmental defects and toxicity.

#### Drug assay to determine compound toxicity in *C. elegans*

The *C. elegans* Bristol strain N2 was used to determine the safe concentrations of CLR01. The assay was performed in a 96-well plate format in liquid culture.[8] Each well contained a final volume of 60 µL, comprising 20–25 animals in the egg stage, CLR01 at final concentrations 0.0001, 0.001, 0.01, 0.1, 1, 10, 50 or100 µM, and OP50 bacteria at a final OD at 595 nm of 0.6–0.7. Worms were grown with continuous shaking at 180 rpm at 20 °C (Shel Lab incubator shaker), and the bacteria OD was measured daily for seven days in the microplate reader (NanoQuant Plate™-Tecan). The effect of the compound on *C. elegans* physiology was monitored by the rate at which the *E. coli* food suspension was consumed as a readout for *C. elegans* growth, survival, or fecundity. DMSO 1% (drug vehicle) and DMSO 5% (toxic condition) were used as controls.

#### Chronic and post-symptomatic treatment of *C. elegans* with CLR01

Chronic pre- and post-symptomatic *C. elegans* treatments with CLR01 were performed in 96-well plates in liquid culture. Pre-symptomatic, chronic, treatment plates comprised: 45–50 animals per well in S-medium, CLR01 at the respective concentrations in DMSO 1% [100-0.0001 µM], and OP50 bacteria (previously inactivated by cycles of freeze/thawing) resuspended in S-medium complete to a final OD at 595 nm of 0.9. Treatment was initiated at the egg stage until day four post-hatching. Post-symptomatic treatment with CLR01 was performed as described above except that animals were kept in drug vehicle (DMSO 1%) until day four post-hatching. Starting on day 4, animals were washed daily and incubated with freshly prepared 0.1-µM CLR01 or 1% DMSO as a control. The plates were sealed to prevent evaporation and incubated at 20 °C with shaking at 180 rpm (Shell lab incubator shaker) for the indicated times.

#### *C. elegans* motility assays

We used a method described previously[47] to determine the percentage of animals with motor impairments after treatment with CLR01 at the indicated concentrations. Briefly, ∼10 animals were transferred simultaneously into the middle of a freshly seeded plate equilibrated at 20 °C. Animals remaining inside a 1-cm circle after 1 min were scored as locomotion-defective. In the chronic pre-symptomatic treatment with CLR01, the motor phenotype was assessed on day four post-hatching. A total of 200-250 animals were scored in 4 to 5 independent assays for each condition. In the post-symptomatic treatment with CLR01, animals’ motor phenotype was assessed on day 4 (before treatment initiation), and the impact of CLR01 post-symptomatic treatment was evaluated on days 6 and 8. A total of ∼200 animals were scored in 4 independent assays for each condition. All assays were performed at room temperature (∼20 °C) using synchronized animals grown at the same temperature. All animals were scored at the same chronological age, and the scoring was performed blinded to treatment.

#### *C. elegans* aggregation assays

*In vivo* images of the head region of AT3q130 animals were obtained using confocal microscopy (Olympus FV1000) under a 60× oil objective (NA = 1.35). The animals were transferred to 3% agarose pads and immobilized using 5 mM levamisole. Z-series imaging was acquired for vehicle- and 0.1 µM CLR01-treated animals using the 515-nm laser line for excitation of yellow fluorescent proteins. The pinhole was adjusted to 1.0 Airy unit of optical slice, and a scan was acquired every 0.5 µm along the Z-axis. The obtained images were analyzed using a MeVisLab tool, as described previously[47, 48]. Two parameters were measured: area of aggregates/total area and number of aggregates/total area. The values shown are the mean (normalized to vehicle-treated control). Three independent trials were performed, and 28 and 30 animals were analyzed in the vehicle and CLR01 treatments.

### Mouse SCA3 model

#### Animal housing conditions

Female transgenic SCA3 and WT-littermates (background C57BL/6J) exposed to CLR01/vehicle treatments were housed at weaning in groups of 5-6 animals in filter-topped polysulfone cages 267 × 207 × 140 mm (370 cm^2^ floor area) (Tecniplast, Buguggiate, Italy), with corncob bedding (Scobis Due, Mucedola SRL, Settimo Milanese, Italy) in a conventional animal facility. The animals were kept under regular laboratory housing conditions: an artificial 12 h light/dark cycle (lights on from 8 am to 8 pm), with a relative humidity of 50–60% and an ambient temperature of 21±1 °C. Mice were under a standard diet (4RF25 during the gestation and postnatal periods and 4RF21 after weaning, Mucedola SRL, Settimo Milanese, Italy) and water ad libitum. Sentinel animals in the same housing room were used to monitor the health status of the mice, according to FELASA guidelines, confirming the Specified Pathogens status. Humane endpoints for the experiment were defined as a 20% reduction of the body weight, inability to reach food and water, presence of wounds in the body, or dehydration. Still, they were not needed in practice as the ages tested in this study were within a range in which the animals did not reach these endpoints. Standardized environmental enrichment in our animal facilities (soft paper and creased paper for nesting behavior) was used.

#### CLR01 treatment groups

Animals were distributed among the different treatment groups in a randomized manner: WT vehicle (n = 19), WT CLR01 (n = 13), SCA3 vehicle (n = 20), and SCA3 CLR01 (n = 14). DNA extraction, animal genotyping, and CAG repeat size analyses were performed as described previously.[87] The CAG repeat mean was not different among SCA3 groups (Mean ± SD; [min-max]_CAG_. vehicle: 140 ± 2.56; [135–143]_CAG_; CLR01: 139 ± 3.04; [136–144]_CAG_). CLR01 (1 mg/Kg per day) or vehicle (0.9% saline) was administered subcutaneously, daily, and five times per week from 4 to 22 weeks of age. CLR01 solution was prepared freshly every week in 0.9% saline. The injection volume was 100 μL and was administered by an independent experimenter (not involved in performing behavioral testing). Behavioral analyses were performed during the diurnal period in groups of 5-6 animals per cage of CMVMJD135 hemizygous transgenic mice and WT littermates treated with CLR01 or vehicle.[6] The same experimenter, blind to treatment, always performed the behavioral tests. At the end of the pre-clinical trial, the animals were euthanized according to their final purpose: either by decapitation or exsanguination perfusion with saline or paraformaldehyde (PFA) 4%. In the latter case, the animals were deeply anesthetized with a mixture of 150 mg/kg ketamine hydrochloride and 0.3 mg/kg medetomidine.

### Mouse phenotypic analysis

#### Body weight

All the animals were weighed at four weeks of age before treatment initiation, and their body weight was monitored every two weeks during the entire study.

#### Beam walk balance test

The test was performed as described previously[88]. Animals were trained by the experimenter for three consecutive days using a 12 mm square-shaped, 1-meter-long beam. Each animal had to traverse the beam three times during the learning phase. On day four, animals were tested in two different beams: the training 12 mm-squared and an additional round beam of a 17-mm diameter. Animals were allowed to fail twice during the test, either by turning around or falling. Quantification of the time each animal took to traverse the beams was registered by the experimenter.

#### Motor swimming test

To analyze voluntary locomotion, the mice were trained for two consecutive days, three trials for each animal, to traverse a clear, 100-cm long, water tank to a safe platform at the end. The latency to cross the water tank was measured from a distance of 60 cm. The water temperature was monitored and set at 23 °C using a thermostat.[88]

#### Spontaneous activity

The mice were transferred to a 15-labelled-square arena (55 × 33 × 18 cm). The number of squares traveled in the arena for 1 minute was counted. The experimenters, blinded to treatment, registered the gait quality as normal or abnormal.

#### Neuropathology analysis

SCA3 and WT littermate mice were deeply anesthetized and transcardially perfused with PBS, followed by PFA 4% in PBS. Spinal cords were harvested and post-fixed overnight in a fixative solution. The following day, spinal cords were transferred to a sucrose 30% solution and further sliced in a vibratome. 40-µm-thick, free-floating sections were stained with cresyl violet or processed for immunohistochemistry with mouse anti-Atx3 (1:500, Millipore, 1H9, Cat #MAB5360). Atx3 positive inclusions, pyknotic cells, and motor neurons were quantified in the ventral horn (100% and 50% of coverage, respectively; n = 5-6 animals/group; 6 slices/animal) and normalized to the total area using an Olympus BX51 stereological microscope and a Visiopharma integrator system software. The integrated density of amyloid fibrils and fibrillar oligomers obtained by immunostaining with anti-OC antibody was measured in mosaic pictures obtained using a 20× objective in an Olympus LPS Confocal FV3000 microscope. The sum of the pixel intensity values (integrated density) was obtained within the area drawn (gray matter of the spinal cord, using the polygon option) in the Fiji software. The integrated density was then normalized for each corresponding area. Negative controls (no primary antibody added) were performed in all immunostaining experiments. In the case of anti-OC immunohistochemistry, we used brain sections of a mouse model of Tauopathy, THY-Tau22,[89] as a positive control. All the samples were codified immediately after euthanasia using numeric codes, and the quantification was performed blinded to genotype and treatment.

### Statistical analysis

All statistical analysis was performed using the IBM SPSS statistics 23 software (SPSS, RRID:SCR_002865) and GraphPad Prism 8.0.1 software.

#### C. elegans

Continuous variables with normal distributions were analyzed according to Kolmogorov-Smirnov (K-S) test p > 0.05, and with homogeneity of variance assessed by Levene’s test, p > 0.05. When these two parameters were confirmed, groups were compared using one-way ANOVA (factor: treatment) using the Dunnet test for post hoc analysis, and when the comparison was only between two groups, we used an independent-sample t-test. In the toxicity tests, in the comparison between groups, we used a non-linear regression curve fit for sigmoidal curves analyzed using a least squares model with two parameters, IC_50,_ and Hill Slope values. Results are presented as the mean ± SEM.

#### Mice

The experimental unit in this study was a single animal. For the motor phenotype analysis, sample size calculations were performed for each behavioral test, assuming a power of 0.8 and a significance level (p-value) of 0.05. The effect size was calculated to be 50%, using mean and standard deviation values obtained previously for transgenic and control groups for each test.[6, 8] Variance homogeneity was evaluated using Levene’s test. Continuous variables with normal distributions (K-S test p > 0.05) were analyzed using a unidirectional or repeated-measure ANOVA. Appropriate post-hoc tests were used to follow up on any significant effects of genotype, treatment, or interaction effects found in the ANOVA having in consideration the unequal sample size of the experimental groups. Behavioral data were subjected to non-parametric Mann-Whitney U-test or Kruskal-Wallis H-tests when variables were non-continuous or when continuous variables did not present a normal distribution (Shapiro-Wilk test p < 0.05). Outliers were considered when the value obtained was 1.5×IQR. A critical value for significance of p < 0.05 was used throughout the study. Statistical reports of all the data analyzed can be found in **Supporting Information Tables S5-S9**.

## ETHICS STATEMENT

All procedures followed European regulations (European Union Directive 2010/63/EU). Animal facilities and the people directly involved in animal experiments (SDS, ANC, PM) were certified by the Portuguese regulatory entity — Direcção Geral de Alimentação e Veterinária. All protocols performed were approved by the Animal Ethics Committees of the Life and Health Sciences Research Institute, University of Minho, and the Center for Neurosciences and Cell Biology, University of Coimbra.

## COMPETING INTERESTS

The authors declare that they have no competing interests.

## FUNDING

This work has been supported by the Portuguese Foundation for Science and Technology (FCT, Portugal) projects POCI-01-0145-FEDER-031173 (PTDC/BIA-BFS/31173/2017), POCI-01-0145-FEDER-007274, UIDB/50026/2020 and UIDP/50026/2020; by the Horizon2020 project PhasAGE (GA 952334). Additionally, this project was supported by the National Ataxia Foundation (NAF Young Investigator – SCA Award attributed to AS and NAF Seed Money research grant attributed to SDS and PM) and by Amyloidosis Foundation (research grant attributed to AS). ESG was supported by the Deutsche Forschungsgemeinschaft (DFG) under Germanýs Excellence Strategy – EXC 2033 – 390677874 and by the instrumentation programme of the DFG (Project-ID 436586093) as well as the CRC1430 (Project-ID 424228829). GB was supported by NIH grant R01AG050721. JAL acknowledges support from the US National Institutes of Health (R35GM145286, S10OD018504) and the US Department of Energy (DE-FC02-02ER63421). SMR acknowledges support from Ataxia-UK, AISA (Associazione Italiana per la lotta alleSindromi Atassiche), ACAH (Catalan Association of Hereditary Ataxias), Swedish SCA-network, and Plata-forma R+SCAs.

## ACKNOWLEDGMENTS

We acknowledge the i3S Biochemical and Biophysical Technologies Platform; and the i3S Advanced Light Microscopy Platform, the i3S Histology and Electron Microscopy Platform, and the ICVS Scientific Microscopy Platform, members of the national infrastructure PPBI - Portuguese Platform of Bioimaging (PPBI-POCI-01-0145-FEDER-022122); FF, DVC, DMF, and ANC were supported by FCT individual fellowships: SFRH/BD/133009/2017, SFRH/BD/147826/2019, SFRH/BD/147947/2019 and SFRH/BPD/118779/2016, respectively. SDS and PMM were awarded contracts from the Scientific Employment Stimulus program from FCT, CEECIND/00685/2020, and CEECIND/03750/2017, respectively.

## Notes

### Competing Interest Statement

The authors have declared no competing interest.

