## Supporting Information for "Allosteric Modulation of Pathological Ataxin-3 Aggregation: A Path to Spinocerebellar Ataxia Type-3 Therapies"

### 1. SUPPORTING METHODS

#### 1.1.Molecular modeling studies of JD:CLR01 Complexes

##### Computational details

The initial coordinates for the GaMD simulations[1] of the JD in the absence and presence of CLR01 molecules were obtained from the first model of the NMR structures with PDB ID: 1YZB[2] and 2AGA[3] (**Figure S2**). The CHARMM-GUI server[4] was employed for the preparation of all systems, generation of their topologies for the CHARMM36m force field[5] and the inputs for AMBER.[6] The protonation states for the titrable protein residues were set at pH 7. The systems were solvated in a rectangular box of TIP3P[7, 8] water molecules with a minimum distance of 15 Å between the protein and the edge of the box. Counterions of NaCl were added for net charge neutralization. The same protocol was applied to study the JD in the presence of CLR01. The JD has in total six lysine and eight arginine residues. In the 1YZB model, two lysine residues (K125, K166) and three arginine residues (R47, R110, and R124) were sterically inaccessible for complex formation with CLR01. In the 2AGA model, two arginine residues (R110 and R124) and one lysine residue (K125) were inaccessible. CLR01 molecules were placed around the remaining lysine and arginine residues, resulting in 1:9 and 1:11 (1YZB or 2AGA): CLR01 complexes, respectively (**Figure S2, right**). Additionally, we prepared a system comprising the JD open model and a single CLR01 molecule positioned at K128. The complexes were simulated following an analogous procedure as for the JD models alone. For CLR01, CHARMM general force field (CGenFF) parameters,[9] tested and used in previous studies,[10, 11] were employed. The particle mesh Ewald (PME) method was used for long-range electrostatic interactions[12] and a cutoff of 12Å for the vdW contributions as implemented in CHARMM-GUI. Each system was subjected to 5000 minimization steps, followed by NVT of 125 ps applying harmonic positional restraints and NPT simulation of 250 ps at 303.15 K and 1 bar of external pressure using Berendsen barostat. Then, 2.4 ns of classical MD were carried out, followed by 8-12 ns of GaMD equilibration until convergence of k0D and k0P to 1. Unrestrained GaMD productions were carried out (3 replicas of 200 ns each) for all the setups using AMBER 20.[13] In the case of the additional system of the JD open in complex with a single CLR01 molecule positioned at K128, the replicas were run for 500 ns of simulation time. The simulations of the JD open structure without the tweezer were extended accordingly. For the analyses of the trajectories and visualization we employed Visual Molecular Dynamics VMD[14] and PyMOL[15] software.

##### Gibbs Energy Calculations

The CL-FEP approach[16] was employed to investigate the binding of the JD to CLR01. The sampling was performed with independent MD simulations on subsystems that included the ligand (CLR01), the host protein (JD), the complex (JD+CLR01), and the bulk solvent (water box). A harmonic wall potential of 50 kcal/mol between the center of mass (COM) of CLR01 and 2 Å distance of the COM of its binding site in the host was applied. An RMSD restraint of 100 kcal/mol was imposed in the simulation setups of the complex and host. Every subsystem was solvated with water molecules using the TIP3P model.[7] The PME method[12] was employed to account for long-range electrostatic interactions and $Na^+$ and $Cl^-$ ions were added for neutralization. The initial minimization simulation was carried out for 10,000 steps, followed by NVT equilibration of 75 000 steps. Subsequently, NPT simulations of 250,000 steps and 2fs timestep using harmonic constraints were performed. Nine individual setups (complex, host, ligand, solvent) corresponding to the nine sterically accessible Lys/Arg were generated for the 1YZB model. We carried out three production replicas of 100 ns each for all setups.[17] The free energy estimation over the 300 ns of sampling was obtained using the second-order cumulant estimator. The analysis was performed with 15 checkpoints and replicated 30 times. The average free energy values among converged checkpoints with further subtraction of the correction factor provided the final estimation of free energy changes.[16]

##### 1.2.Numerical Methods for Biophysical Model of Atx3 Self-Assembly

DLS data measured during Atx3 13Q are well described by a nucleation-and-growth model of phase separation.[18] The main simplifying hypothesis adopted during the derivation of this model are: (i) primary nucleation is described by a second-order rate equation on supersaturation, (ii) growth and secondary nucleation are autocatalytic processes whose rates increase with the mass concentration of particles and are first-order functions of supersaturation, (iii) no significant occurrence of breakage/fragmentation/ coalescence events; (iv) the volume of the supersaturated solution remains approximately constant; (v) no surface tension effects are considered.[19] Under these conditions, the time evolutions of mass $M(t)$ and mean size $N(t)$ of new particles formed by NAG are given by the equations:[18, 19]

$$\alpha(t) = 1 - \frac{1}{k_b[\exp(k_\alpha \Delta\mu_0 t) - 1] + 1} \tag{1a}$$

$$\frac{N_1}{N(t)} = \frac{k_b}{1-k_b}\left[\frac{\ln\big(1-\alpha(t)\big) + k_\alpha \Delta\mu_0 t}{\alpha(t)(1-k_b)}\left(1 - \frac{k_2}{k_\alpha}\frac{N_1}{N_2}\right) - \left(1 - \frac{k_2}{k_\alpha k_b}\frac{N_1}{N_2}\right)\right] \tag{1b}$$

where $\alpha(t) = M(t)/M_\infty$ is the reaction conversion and $M_\infty$ is the total mass; $N_1$ and $N_2$ are the sizes of primary and secondary nucleation particles, respectively; $\Delta\mu_0$ is the initial supersaturation approximated as $\Delta\mu_0 \approx (c_0 - c_\infty)/c_\infty$, with $c_0$ and $c_\infty$ representing the inital protein concentration and the protein solubility, repectively; and $k_\alpha$ and $k_b$ are functions of the rate constants characterizing the microscopic steps of primary nucleation ($k_n$), surface growth ($k_+$) and secondary nucleation ($k_2$):

$$k_\alpha = (k_+ + k_2) \tag{2a}$$

$$k_b = \frac{k_n}{k_+ + k_2} \quad (2b)$$

The scattering intensities represented by vertical lines in Fig. 2 and Supplementary Fig. S5 are computed assuming the equivalence between $N(t)/N_1$ and $[R(t)/R_1]^3$ ratios, where $R$ refers to hydrodynamic radii. Each particle is modelled as a Mie scatterer, whose scattering intensity is calculated using the MATLAB code *MatScat*.[20] A good agreement between the NAG model and the DLS data in the absence of CLR01 is obtained by adopting a simplified fitting procedure that: (i) assumes $k_2/k_\alpha \approx 1$ when the increase in scattering intensity takes place at nearly invariant $R(t)$; (ii) assumes $k_2/k_\alpha \approx 0$ when both the scattering intensity and $R(t)$ increase over time. In the presence of CLR01 increasing values of the $k_2/k_\alpha$ ratio were tested until an optimal numerical agreement between measured and simulated DLS data was verified. In all cases, the values of $k_\alpha \Delta\mu_0$ and $k_b$ were directly obtained from solving Eqs. 1a and 1b using the half-life coordinates $(\alpha(t_{50}) = 0.5)$ measured during thioflavin-T (ThT) fluorescence experiments. This means that the model parameters that describe the DLS data also describe the kinetic measurables describing ThT progress curves. The values of $k_n\Delta\mu_0$, $k_+\Delta\mu_0$ and $k_2\Delta\mu_0$ were determined from Eqs. 1a and 1b using the known parameters $k_\alpha\Delta\mu_0$ and $k_2/k_\alpha$. While the results in **Fig. 2** and **Figs. S7a, b** support the validity of our theoretical analysis, the size distributions obtained for long aggregation times exhibit features that the NAG model does not predict. One of these features is the formation of Atx3 77Q aggregates with hydrodynamic radii $R_H$ in the micrometre range (**Figure 2c**). To account for this population of large species, an additional step of fibril agglomeration (that does not increase ThT fluorescence) has to be considered only for the Atx3 77Q variant. Another interesting feature is the population of very small oligomers ($R_H$=10 nm) observed after 72h incubation for both variants in the presence of CLR01 (**Figure S7a, b**). This result suggests that CLR01 induces additional secondary pathways such as fibril fragmentation or secondary nucleation by which very small oligomers are generated.

##### 1.3. Photo-Induced Cross-linking of Unmodified Proteins (PICUP) assay

PICUP reactions were performed according to previously described protocols, with minor modifications.[21, 22] Briefly, for each reaction, the purified proteins and CLR01 solutions were mixed diluted to a final concentration of 10 µM in the assay buffer (100 mM sodium phosphate, pH 7.4). For each reaction, the protein and CLR01 solutions were mixed to the final concentrations in 18 µL PCR tubes, followed by the addition of 1 µL Ru(Bpy) and 1 µL APS. The final concentration of protein was 10 µM, the CLR01 concentrations were 1, 3, 10, 30 or 100 µM, the concentration of Ru(Bpy) and APS were 40 µM and 800 µM, respectively. The reaction started with irradiation for 1 s, and quenching immediately with 1 µL DTT. Samples were fractionated by SDS-PAGE using Tris-

Acetate. Non-cross-linked proteins were used as a control. Protein bands were visualized by silver staining (Invitrogen). Gels were scanned and the abundance of oligomeric species in each lane was calculated by densitometry analysis relative to the entire lane using Image J. The data represents the average of 3 replicates.

#### 2. SUPPORTING RESULTS

##### 2.1.GaMD Simulations: Effect of CLR01 on the structure of Josephin Domain (JD) and prediction of CLR01 favored binding sites

We performed Gaussian accelerated molecular dynamics (GaMD) simulations to explore the inhibition mechanism of Atx3 aggregation by CLR01. To investigate the favored binding sites of CLR01 in the JD, we analyzed the non-covalent contacts of the tweezer molecules with the amino acids in the JD over the GaMD. The simulations for open conformation in the presence of CLR01 showed several residues with a large population of contacts (**Table S1**). The estimated binding free energy changes for the complexes between CLR01 and selected residues of JD on the open conformation revealed seven sites in which binding of CLR01 to exposed lysine and arginine residues on the JD is favored (**Figure S4c**). However, R182, K85, R101, K117, R59, and R103, also showed a high frequency of contact and a calculated binding free energy compatible with a putative interaction, no chemical shift perturbations were observed in the NMR data. Previous data has shown that, although arginine residues may be favored electrostatically, lysine residues are sterically more accessible and have consistently shown to have a higher affinity for CLR01,[23, 24] in full agreement with our simulations for JD (**Figure S4b**). In fact, the smaller and less bulky side chain of lysine can thread more easily than arginine into the tweezer's cavity without inducing structural distortions of the torus-shaped arrangement of alternating benzene and norbornadiene rings.[25] Concerning the residues with thermodynamically favored CLR01 interactions, K128 emerges as the preferred CLR01 binding site and had the second highest SASA value for lysine in JD, after K8. K8 and R182, located at the N- and C-terminus, respectively, were involved in hydrogen bonds with E7 (12.9%) and E10 (14.8%) for the former and with Q129 (13.9%) and D145 (15.0%) for the latter, potentially hindering stable CLR01 binding. For K85, which exhibited a similar SASA value to K128, the occupancy of the hydrogen bond with E90 was 29.1%. The simulations for open conformation in the presence of CLR01 at K128 showed several residues with a large population of contacts close to the termini of the JD, including hydrophobic cluster (L178, M180, I181) that may further stabilize complex formation at this site.

#### 2.2. PICUP Assays and SEC analysis of Atx3 oligomerization

To assess the effect of CLR01 on the initial stages of Atx3 assembly, we used the Photo-Induced Cross-linking of Unmodified Proteins (PICUP) assay.[21, 22] Cross-linked dimers of the JD, and a range of Atx3 13Q and Atx3 77Q oligomers, including putative dimers, trimers, and tetramers, could be visualized in the PICUP assay (**Figure S6b**). CLR01 weakly decreased the relative abundance of oligomeric species for both Atx3 variants and for the JD. The $IC_{50}$ values were between 170 to 200 μM), corresponding to a 17- to 20-fold molar ratio of the molecular tweezer (**Figure S6b**). The effect of CLR01 on the overall decrease in Atx3 oligomerization was further corroborated by the analysis of the time-dependent distribution of monomer and high molecular weight (HMW) soluble species by size-exclusion chromatography (SEC). CLR01 caused a delay in the conversion of monomers to HMW species for both Atx3 variants (**Figure S6c**). This delay was analyzed quantitatively by comparing the relative peak areas of HMW oligomers and Atx3 monomers. In the presence of CLR01, at the end-point of the aggregation assay (140h), ~50% Atx3 13Q remained in its monomeric form, whereas in the control condition, less than 20% of the monomeric species remained after 90h at 37 °C. Similarly, most Atx3 77Q monomers were converted to HMW species after 72 h, while monomers were still detectable even after 90 h of incubation in the presence of CLR01. This effect was not observed by the incubation with the control molecule, CLR03.

### 3. SUPPORTING FIGURES AND CAPTIONS

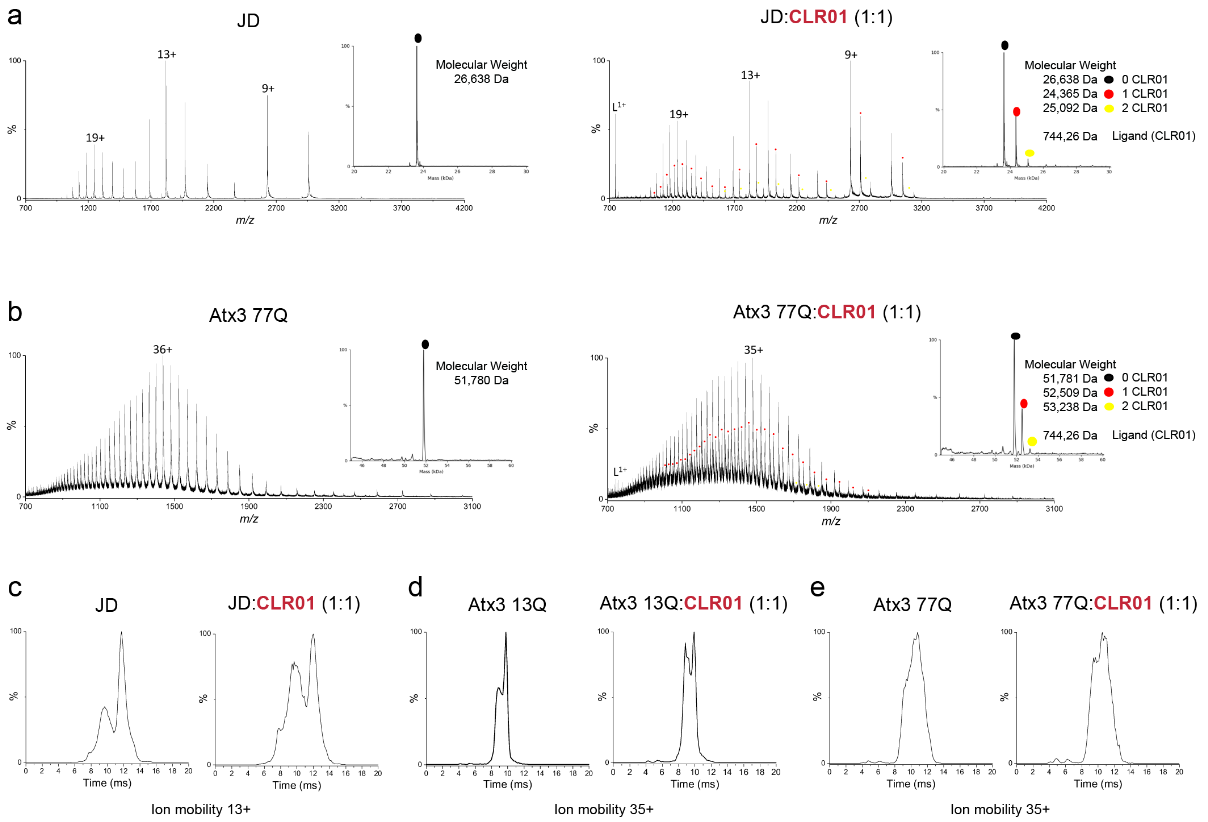

**Figure S1 – CLR01 forms a complex with the isolated JD and with full-length Atx3**. **a)** ESI mass spectra of JD alone (left) and in complex with CLR01 (right). **b)** ESI mass spectra of Atx3 77Q alone (left) and in complex with CLR01 (right). The insets in a) and b) show the deconvoluted spectra and highlight the two CLR01 binding events. **c)** IM-MS of the 13+ charge state of JD in the absence of presence of CLR01. **d)** IM-MS of the 35+ charge state of Atx3 13Q in the absence of presence of CLR01. **e)** IM-MS of the 35+ charge state of Atx3 77Q in the absence of presence of CLR01.

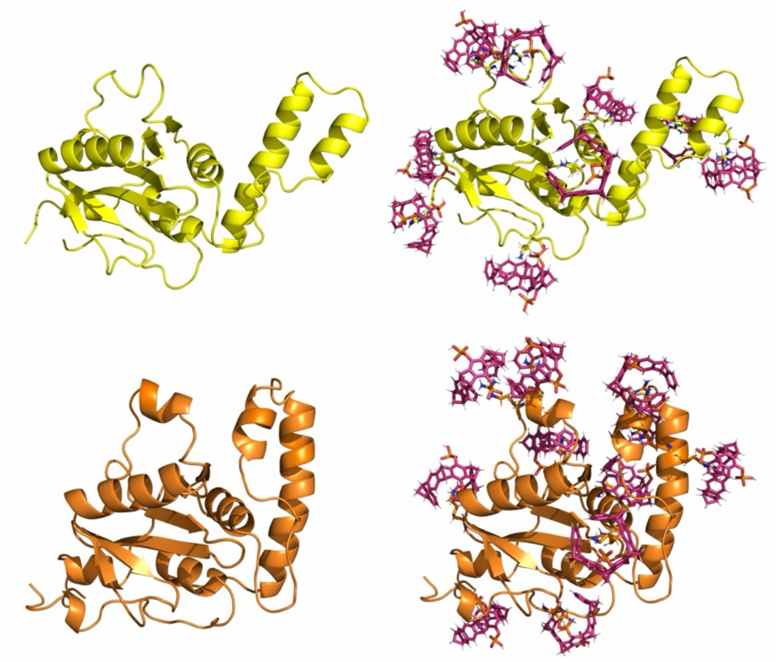

**Figure S2 -** Initial structures of the JD domain used for the simulations: 1YZB model (open, in yellow) and 2AGA model (closed, in orange) in the absence (left) and presence (right) of CLR01 molecules. Lysine and arginine residues as well as CLR01 are shown in licorice representation.

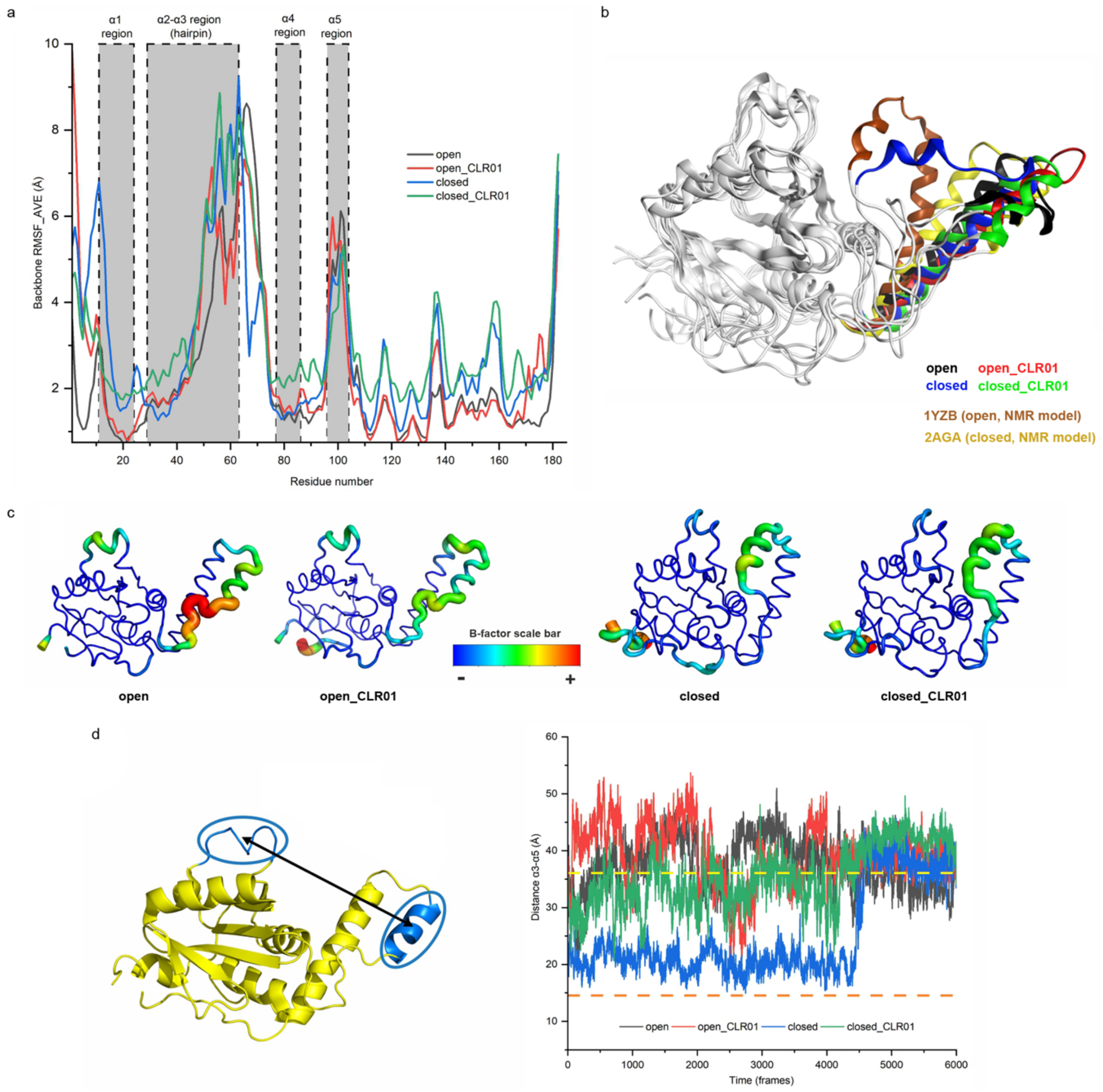

**Figure S3 - Conformational variability of JD throughout 600 ns of GaMD simulations in absence or presence of CLR01 molecules**. **a)** Root Mean Square fluctuation (RMSF) plot. Each point represents the mean fluctuation per residue calculated over the complete GaMD trajectory. The helixes α1, α2, α3, α4 and α5 are highlighted in grey and dashed boxes. The most dynamic region is the hairpin loop (α2 and α3, S29-Q63), which is responsible for the large-scale conformational change observed in the simulations ($RMSF_{hairpin}$ > 7Å. **b)** Superposition of representative structures of the most populated cluster for each simulated system (JD open, JD open_CLR01, JD closed, and JD closed_CLR01). The NMR models 1YZB and 2AGA were included as reference for JD's open and closed states, respectively. **c)** The average backbone B-factor profiles as function of RMSF were calculated from the GaMD trajectories of JD with or without CLR01. The highest B-factor (highest flexibility) for each structure is coloured in red and the lowest in dark blue. The thickness of the protein backbone is also proportional to the B-factors. **d)** Distance between the centre of mass of the α3 and α5 regions of the JD in the absence or presence of CLR01. The reference distance values, taken from the NMR models, are represented as two horizontal dash lines for the open (yellow) and the closed (orange) states of the JD. The analysis reveals the inherent flexibility of the JD, exhibiting a coexistence of both states, whereas the JD open_CLR01 and JD closed_CLR01 systems undergo partial or complete transitions to a half-open conformation upon the addition of CLR01. The closed system, in the absence of CLR01 (closed, blue graph), transitioned to the open configuration in one of the GaMD replicas, in agreement with previous reports of the open form as the favored state of the JD.[26]

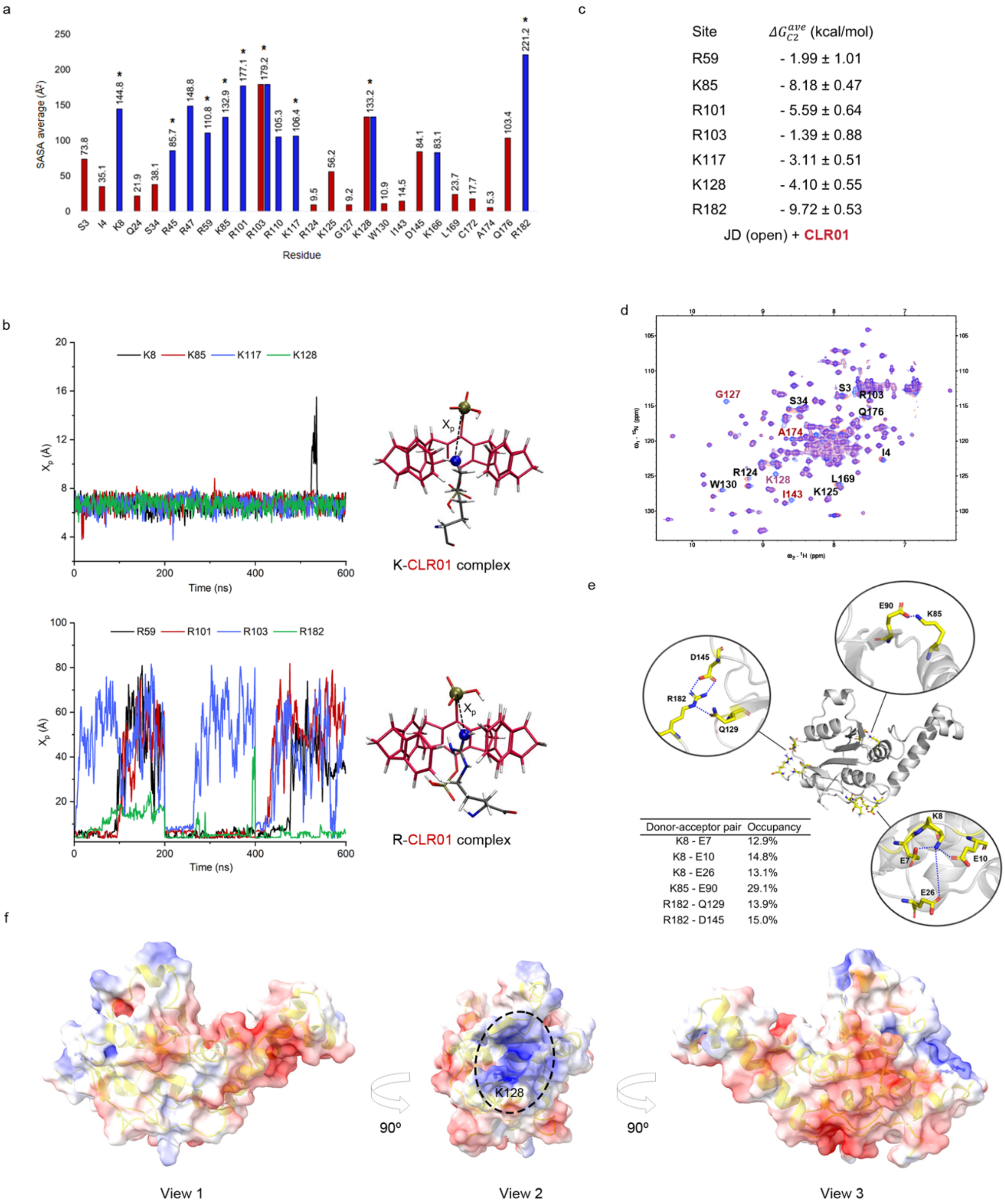

**Figure S4 – Computational and experimental analysis of CLR01 interaction with JD**. **a)** SASA analysis of all JD (open conformation) R/K residues (blue); residues for which chemical shifts were observed in NMR (panel d) are shown in red. The values on top of the bars show the average SASA for each residue computed over the three GaMD trajectories of JD; the star symbol highlights the R/K residues simulated in the complex with CLR01. **b)** Plot of the $X_p$ geometric parameter corresponding to the distance between the closest nitrogen atom of K/R residues side chain and the phosphorus atom in CLR01. **c)** Binding Gibbs energies (calculated using CL-FEP with the second order cumulant (C2) estimator) for the R/K residues in complex with CLR01 for which binding was predicted to be thermodynamically favourable. **d)** Overlay of SOFAST-HMQC spectrum of JD alone (100 uM; blue) and upon addition of a twofold molar excess of CLR01 (red), recorded at 21°C on a 800 MHz spectrometer. Residue numbers identify resonances with the largest CSPs. **e)** Hydrogen bond interactions of K8, K85 and R182, illustrating their most frequent interactions over the simulations. **f)** Electrostatic potential surface of a representative structure from the most populated cluster of JD (open conformation) generated with APBS.[27] In view 2, the area with the highest positive charge density, localized around residue K128 (in balls and sticks), is highlighted.

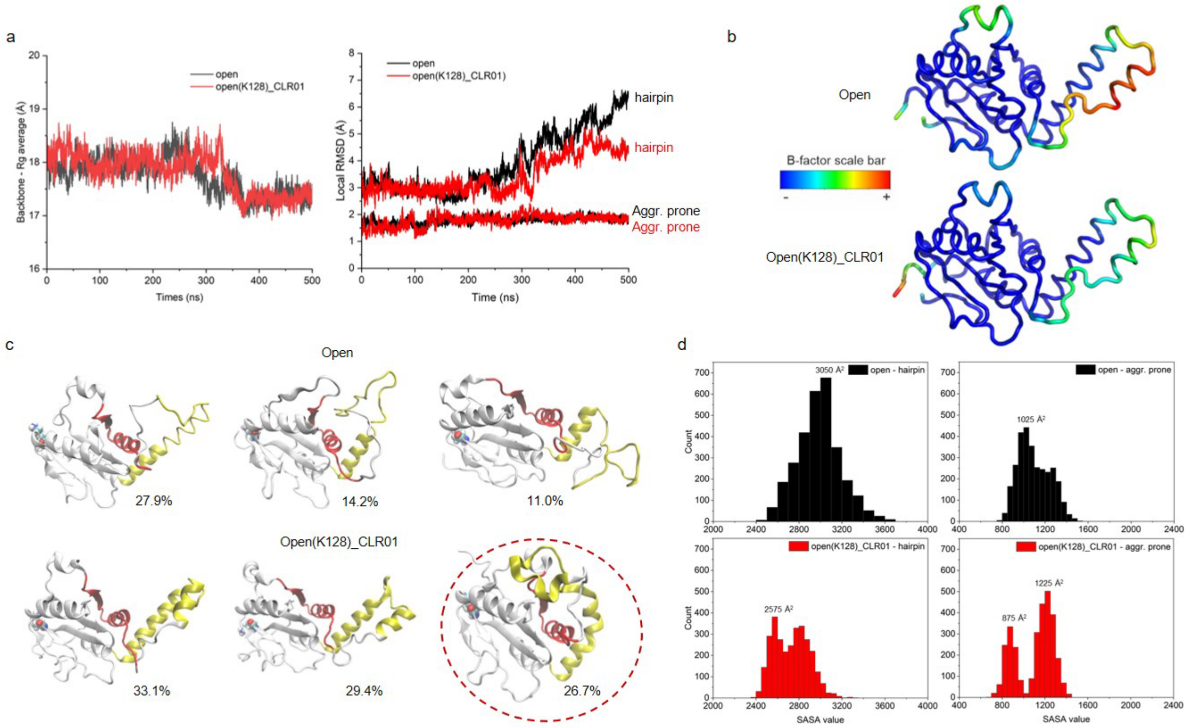

**Figure S5 – CLR01 binding to K128 induces an allosteric effect on the JD helical hairpin. a)** Radius of gyration (Rg) of the protein backbone and root-mean-square deviation (RMSD) focused on the helical hairpin and aggregation-prone region. Both analyses represent average values from three GaMD production replicas of 500 ns each and were obtained from simulations of the JD alone or in complex with CLR01 bound to K128. **b)** Average backbone B-factor profiles as a function of root-mean-square fluctuation (RMSF) were calculated from the GaMD trajectories (3x500 ns) for JD alone or in complex with CLR01. The highest B-factors (indicating the largest flexibility) are colored red, while the lowest are dark blue. **c)** Clustering analysis of the last 100 ns from each GaMD replica (3x100 ns) for the JD alone or in complex with CLR01. The centroids of the three most populated clusters of structures (with corresponding population percentages) reveal the high flexibility of the helical hairpin motif. In the JD:CLR01 complex, a state (dashed circle) was identified where the hairpin moves upon the aggregation-prone region, corresponding to a population of 26.7%. **d)** Histograms of solvent-accessible surface area (SASA) values focused on the hairpin and aggregation-prone region over the last 100 ns from each GaMD replica (3x100 ns) for JD or JD:CLR01. The labels on the histogram peaks represent the values with the highest frequency in each case.

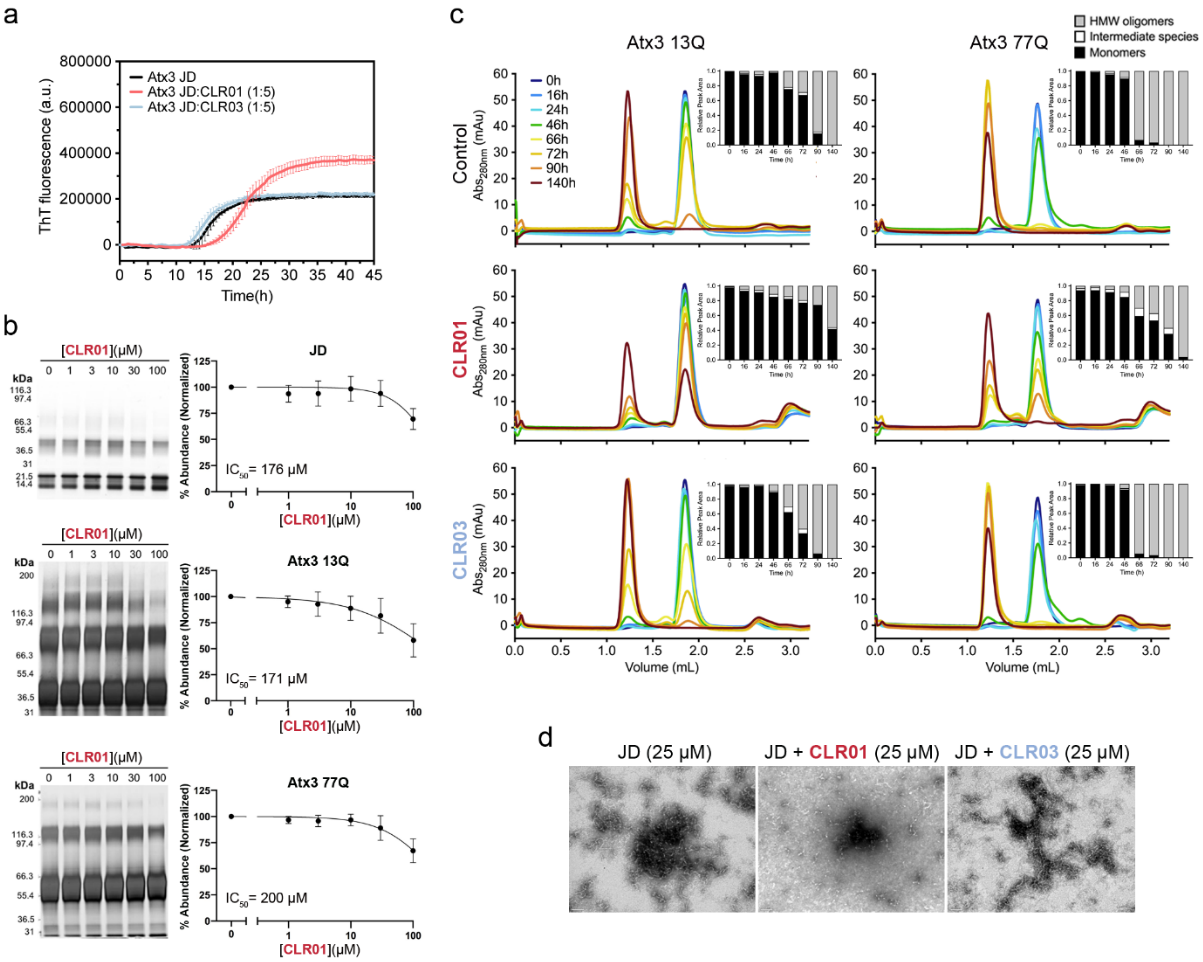

**Figure S6 - CLR01 delays the self-assembly of JD and full-length Atx3. a**) ThT measurement of JD (5 µM) amyloid self-assembly kinetics in the absence and presence of 5-fold molar excess of CLR01 or CLR03. Error bars correspond to SD of 3 replicates. **b**) Photo-induced cross-linking of unmodified proteins (PICUP) assay of 10 µM JD, Atx3 13Q, or Atx3 77Q in the presence of 0, 1, 3, 10, 30 or 100 µM of CLR01, followed by SDS-PAGE analysis, to determine CLR01 half-maximal inhibitory concentration ($IC_{50}$) of Atx3 isoforms oligomerization. Error bars correspond to SD of 3 replicates. **c**) Time-course distribution of soluble monomeric and oligomeric species of 5 µM Atx3 13Q or Atx3 77Q in the absence or presence of 25 µM CLR01 or CLR03 monitored by analytical SEC. Relative peak area (RPA) graphical representation of each time point is presented next to the corresponding SEC profile. **d**) Analysis of JD (5 µM) fibrillar structures in the presence of CLR01 (25 µM) or CLR03 (25 µM) at the endpoint of the aggregation assay, monitored by negative staining TEM. Scale bars correspond to 100 nm.

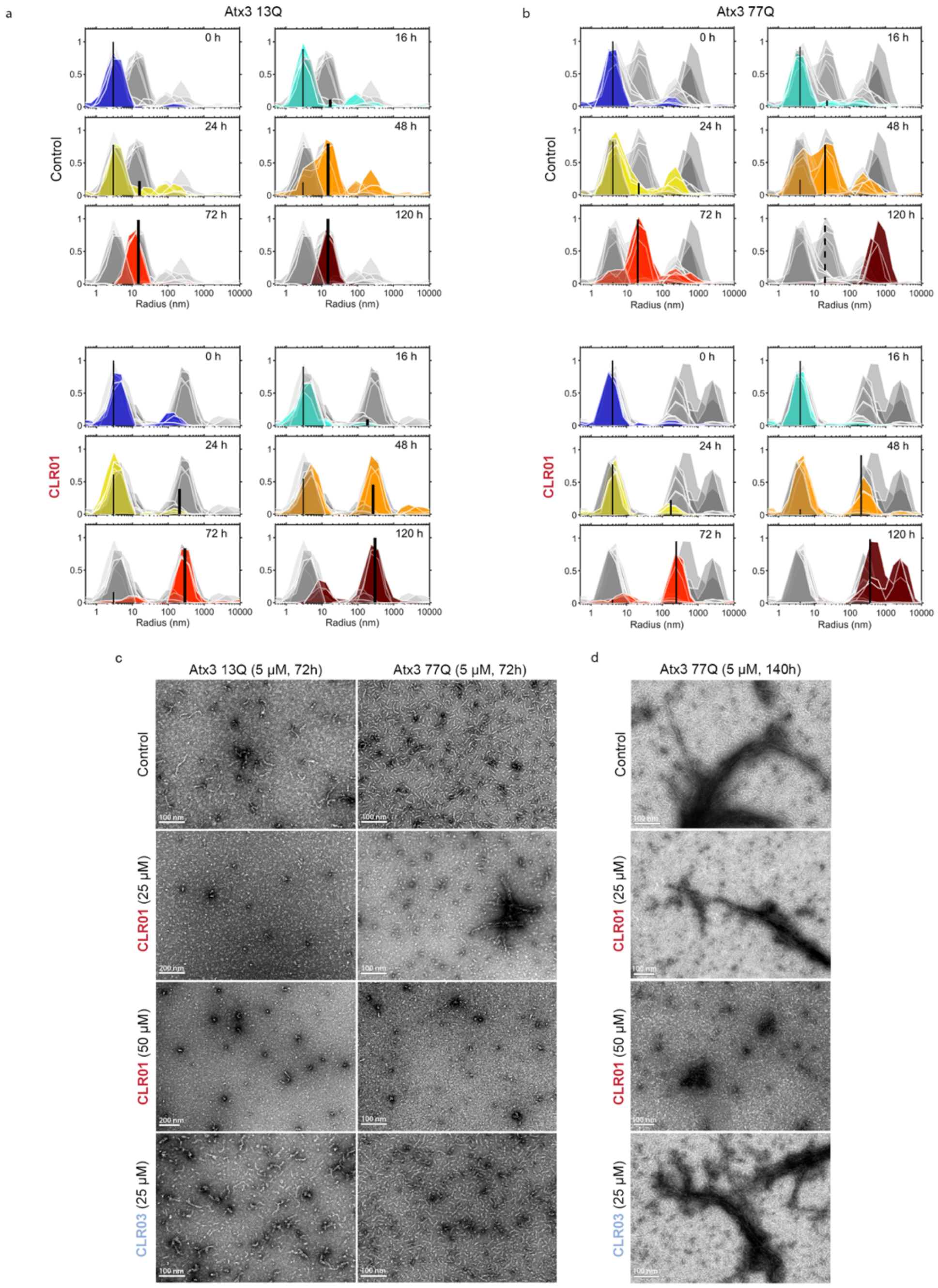

**Figure S7 – CLR01 binding modifies the Atx3 aggregation pathway.** Dynamic light scattering (DLS) intensity (a.u.) distribution at 0, 16, 24, 48, 72 or 120 h of aggregation of 5 µM Atx3 13Q (**a**) or Atx3 77Q (**b**) alone or in the presence of 25 µM CLR01 at 37 °C. Different colors show different time points. The measured size distributions are represented separately for each time point, with the remaining time points represented in greyscale. Vertical lines: scattering intensities predicted by a nucleation-and-growth model for the population of monomers (thinner lines) and the population of amyloid-like fibrils (thicker lines); solid or dashed lines show good or bad agreements between model predictions and measured data. **c**) TEM analysis of the aggregate species formed by wild-type or mutant Atx3 after 72 h of incubation at 37 °C, in the absence or presence of CLR01 or CLR03. In the presence of 25 µM CLR01, a minor population of fibrillar agglomerates were observed for Atx3 77Q. **d**) TEM analysis of the aggregate species formed at the endpoint of the aggregation assay of Atx3 77Q after 72 h of incubation at 37 °C in the absence or presence of CLR01 or CLR03. Scale bars correspond to 200 nm or 100 nm, as indicated in the images.

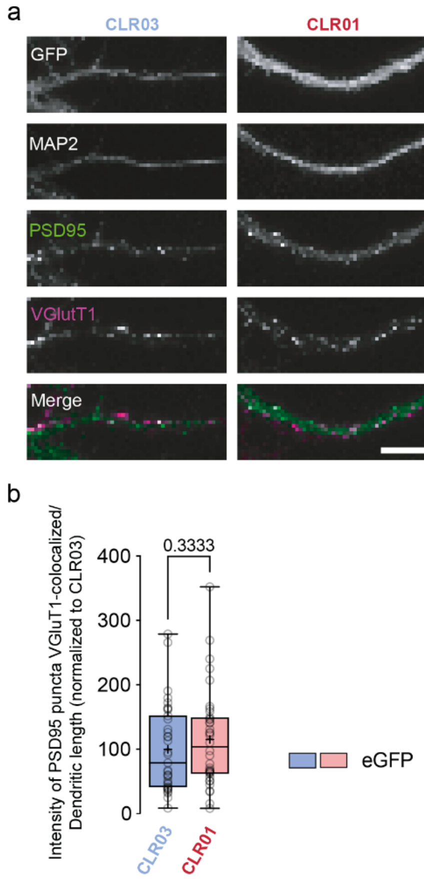

**Figure S8 - CLR01 does not affect glutamatergic synaptic clusters in cortical cells. a**) Immunolabelling of MAP2, PSD95 and VGluT1 in cortical neurons transfected with a plasmid encoding eGFP and incubated with CLR03 or CLR01. **b**) Fluorescence intensity of PSD95/vGLUT1 clusters was quantified (N=3 independent preparations per experiment, n=35-36 neurons per condition). Statistical analyses were performed using a t-test. Boxes show 25th and 75th percentiles, whiskers range from the minimum to the maximum values, and the horizontal line shows the median value. Mean is represented by "+".

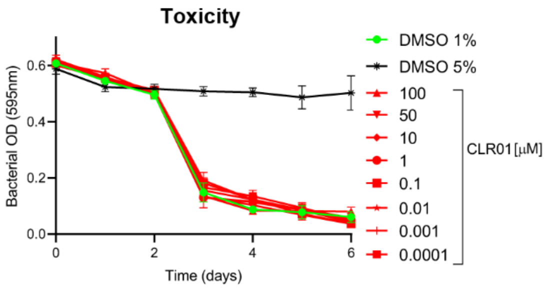

**Figure S9 - Treatment of WT *C. elegans* with CLR01 compound did not affect animals' survival, development and fecundity**. Graphs represent daily bacterial consumption measured by optical density (OD) at 595 nm of animals treated with CLR01, DMSO 1% (negative control) and 5% (positive control) were used as non-toxic and toxic treatment controls, respectively. For each condition tested, the mean OD was calculated for each day from three replicates and plotted over time. The highest safe concentration defined for CLR01 compound was of 100 μM, as the OD decrease paralleled the drug vehicle control samples. Raw data were statistically compared using a non-linear regression (curve fit) for sigmoidal curves and analyzed using a least squares model with 2 parameters ($IC_{50}$ and Hill Slope values).

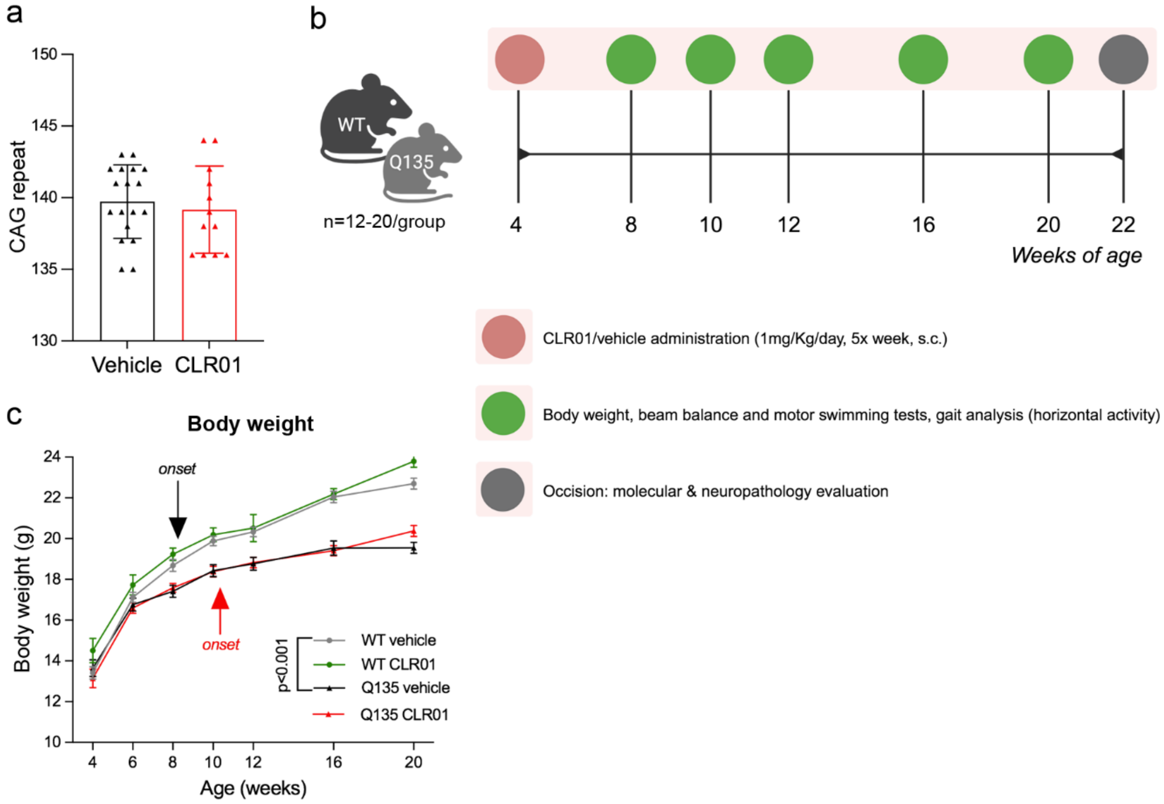

**Figure S10 - Detailed description of the experimental groups used in the pre-clinical trial using CLR01.** **a**) The CAG repeat length distribution was not different between vehicle- and CLR01-treated mice. **b**) Graphical representation of experiment timeline and behavioral testing. **c**) CLR01 treatment did not affect animals' body weight, both in SCA3 and WT mice.

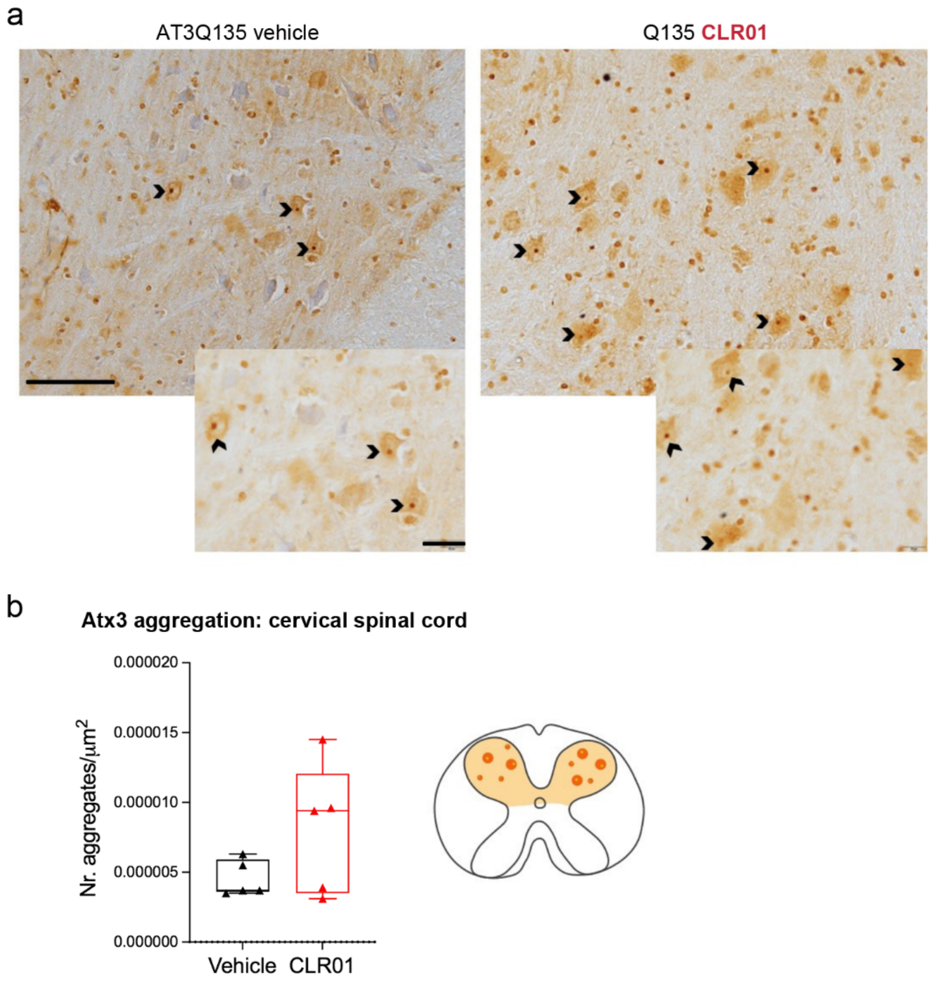

**Figure S11 - Effect of CLR01 chronic administration on Atx3 aggregation in the cervical spinal cord of SCA3 mice. a)** Representative images for Atx3 immunohistochemistry reveal intranuclear inclusions in the cervical spinal cord. **b)** Quantification of these end-stage nuclear Atx3-positive aggregates showed no statistically significant differences upon treatment with CLR01; nevertheless, a trend towards an increase by CLR01 treatment is supported by the large effect size observed (Cohen'd $d = 0.832$). Scale bars represent 100 µm for the lower magnification images and 20 µm for the higher magnification images. Sample size and statistical details are shown in **Supplementary Information Table S9**.

#### 4. SUPPORTING TABLES

**Table S1.** CLR01 K128 binding site. Estimated fraction of contacts (%) between the JD sequence and CLR01 molecule at a 5Å distance over the GaMD trajectories (three replicas taken together). The results for the simulations of the JD in the open conformation in complex with nine CLR01 molecules on selected R/K residues and for one CLR01 molecule at K128 are reported. Amino acids that displayed chemical shifts in the NMR HSQC spectra upon CLR01 binding to the JD are highlighted in red.

| JD(open) + 9CLR01 simulations | | JD(open) + CLR01 on K128 simulations | |
|---|---|---|---|
| Residue | Fraction | Residue | Fraction |
| M1 | 0.42% | M1 | 3.08% |
| E2 | 0.03% | E2 | 0.25% |
| I107 | 21.03% | P106 | 0.25% |
| N108 | 30.28% | I107 | 34.73% |
| E109 | 5.58% | N108 | 38.60% |
| R110 | 40.10% | E109 | 10.55% |
| **K125** | **87.18%** | R110 | 33.63% |
| L126 | 0.85% | R124 | 0.03% |
| **G127** | **83.95%** | **K125** | **78.30%** |
| **K128*** | **100.00%** | L126 | 0.47% |
| Q129 | 19.35% | **G127** | **80.48%** |
| **W130** | **0.45%** | **K128*** | **100.00%** |
| F131 | 0.17% | Q129 | 16.00% |
| L142 | 0.45% | **W130** | **0.80%** |
| **D145** | **28.52%** | F131 | 0.02% |
| T146 | 0.05% | L142 | 0.03% |
| D168 | 0.63% | S144 | 0.13% |
| D171 | 6.85% | **D145** | **15.68%** |
| **C172** | **1.27%** | T146 | 0.02% |
| E173 | 9.28% | D168 | 0.13% |
| **A174** | **3.37%** | D171 | 1.00% |
| D175 | 8.63% | **C172** | **0.15%** |
| **Q176** | **1.73%** | E173 | 0.70% |
| L177 | 2.25% | **A174** | **2.80%** |
| L178 | 21.93% | D175 | 50.78% |
| Q179 | 78.77% | **Q176** | **2.30%** |
| M180 | 56.58% | L177 | 30.33% |
| I181 | 33.17% | L178 | 39.48% |
| R182 | 43.52% | Q179 | 24.00% |
| | | M180 | 27.10% |
| | | I181 | 36.27% |
| | | R182 | 17.88% |

(*): Center of the binding site.

**Table S2** - Report of statistical analysis performed in primary cortical neurons expressing the expanded (eGFP-Atx3 84Q) and non-expanded (eGFP-Atx3 28Q) forms of Ataxin 3. Comparisons between conditions were carried out using Kruskal-Wallis Test followed by Dunn's multiple comparisons test.

| Figure | Treatment | Transfection | Mean percentage of PSD95/vGluT1 puncta intensity ± SD | Statistical report | Sample size (n) |
|---|---|---|---|---|---|
| Fig. 4a-b. (Puncta intensity as a % of control) | N/A | eGFP | 100 ± 78.37 | | 3 (33cells) |
| | | eGFP-Atx3 28Q | 105.1 ± 83.63 | eGFP vs. eGFP-atx3 28Q ( p>0.9999) | 3 (36 cells) |
| | | eGFP-Atx3 84Q | 63.60 ± 53.88 | eGFP-atx3 28Q vs. eGFP-atx3 84Q (p=0.0498)<br>eGFP-atx3 vs. eGFP-atx3 84Q (p=0.0912) | 3 (36 cells) |

**Table S3** - Report of statistical analysis performed in primary cortical neurons treated with CLR01 and CLR03. Comparisons between conditions for each compound were carried out using Kruskal-Wallis Test followed by Dunn's multiple comparisons test.

| Figure | Treatment | Transfection | Mean percentage of PSD95/vGluT1 puncta intensity ± SD | Statistical report | Sample size (n) |
|---|---|---|---|---|---|
| Fig. 4c-d. (Puncta intensity as a % of control) | CLR03 | eGFP | 100 ± 67.8 | | 3 (35cells |
| | | eGFP-Atx3 28Q | 75.9 ± 77.3 | eGFP vs. eGFP-atx3 28Q (p=0.612) | 3 (41 cells) |
| | | eGFP-Atx3 84Q | 58.9 ± 58.28 | eGFP vs. eGFP-atx3 84Q (p=0.0048)<br>eGFP 28Q vs. eGFP-atx3 84Q (p>0.9999) | 3 (40 cells) |
| | CLR01 | eGFP | 100 ± 63.2 | | 3 (36 cells) |
| | | eGFP-Atx3 28Q | 97.1 ± 66.7 | eGFP vs. eGFP-atx3 28Q (p>0.9999) | 3 (39 cells) |
| | | eGFP-Atx3 84Q | 72.7 ± 44.1 | eGFP vs. eGFP-atx3 84Q (p=0.2081)<br>eGFP 28Q vs. eGFP-atx3 84Q (p=0.3847) | 3 (36 cells) |

**Table S4** - Statistical report of statistical analysis performed in primary cortical neurons treated with CLR01 and CLR03. Comparisons between each compound were carried out using Two-way ANOVA followed by Sidak's test for multiple comparisons.

| **Figure** | **Treatment** | **Transfection** | **Mean percentage of neurons with aggregate ± SD** | **Statistical report** | **Sample size (n)** |
|---|---|---|---|---|---|
| Fig. 4e,f. (% of cell with aggregates) | CLR03 | eGFP | 0 ± 0 | Interaction: $F_{(2,12)}$ = 0.1227, p = 0.8856, $\eta^2_p$ = 0.2708<br>Treatment: $F_{(1,12)}$ = 0.06825, p = 0.7983, $\eta^2_p$ = 0.0753<br>Transfection: $F_{(2,12)}$ = 39.15, p < 0.0001, $\eta^2_p$ = 83.41 | 3 (31cells) |
| | | eGFP-Atx3 28Q | 31.2 ± 13.7 | | 3 (32 cells) |
| | | eGFP-Atx3 84Q | 54.9 ± 5.0 | | 3 (33 cells) |
| | CLR01 | eGFP | 0 ± 0 | | 3 (32 cells) |
| | | eGFP-Atx3 28Q | 30.3 ± 22.9 | | 3 (33 cells) |
| | | eGFP-Atx3 84Q | 59.9 ± 4.8 | | 3 (35 cells) |

**Table S5** - Report of statistical analysis performed in primary cortical neurons treated with CLR01 and CLR03. Comparisons between each compound were carried out using Mann-Whitney test.

| **Figure** | **Treatment** | **Transfection** | **Mean percentage of PSD95/vGluT1 puncta intensity ± SD** | **Statistical report** | **Sample size (n)** |
|---|---|---|---|---|---|
| Fig. S8. (Puncta intensity as a % of control) | CLR03 | eGFP | 100 ± 67.7 | p=0.333 | 3 (35cells) |
| | CLR01 | eGFP | 115.1 ± 75.3 | | 3 (36 cells) |

**Table S6 -** Descriptive statistics regarding locomotion defects of AT3q130 animals upon chronic pre-symptomatic treatment with CLR01. Means of the locomotion defective phenotype are reported for all tested concentrations of CLR01. Comparisons between each concentration was carried out using One-way ANOVA followed by a bilateral Dunnett test for multiple comparisons. Efficiency was assessed using the following formula: Efficiency (%) $= \frac{\text{AT3q130}_{\text{DMSO}} - \text{AT3q130}_{\text{drug}}}{\text{AT3q130}_{\text{DMSO}} - \text{WT}}$.
Effect size was given by Cohen's d $= \frac{\text{Mean}_{\text{AT3q130}} - \text{Mean}_{\text{drug}}}{\text{Pooled SD}}$.

| Strain | Concentration (µM) | Mean locomotion defective ± SD | Group comparison (Mean locomotion defective ± SD) | *P*-value | Effect Size | Efficiency (%) |
|---|---|---|---|---|---|---|
| **AT3q130 (AM685)** | 0 (vehicle) | 47.0 ± 8.7 | WT 9.7 ± 4.5 | ***p*<0.001** | | |
| | 0.0001 | 25.9 ± 10.7 | AT3q130 vehicle 46.8 ± 10.1 | ***p*=0.008** | 2.85 | 56 |
| | 0.001 | 27.2 ± 9.2 | AT3q130 vehicle 46.8 ± 10.1 | ***p*=0.014** | 2.90 | 53 |
| | 0.01 | 22.4 ± 5.0 | AT3q130 vehicle 46.8 ± 10.1 | ***p*=0.002** | 4.36 | 66 |
| | 0.1 | 22.1 ± 7.0 | AT3q130 vehicle 46.8 ± 10.1 | ***p*=0.001** | 4.06 | 67 |
| | 1 | 28.6 ± 7.4 | AT3q130 vehicle 46.8 ± 10.1 | ***p*=0.024** | 2.95 | 50 |
| | 10 | 32.2 ± 8.9 | AT3q130 vehicle 46.8 ± 10.1 | *p*=0.100 | 2.20 | 40 |
| | 50 | 30.4 ± 12.4 | AT3q130 vehicle 46.8 ± 10.1 | *p*=0.052 | 2.08 | 45 |
| | 100 | 32.5 ± 7.9 | AT3q130 vehicle 46.8 ± 10.1 | *p*=0.112 | 2.26 | 39 |
| **WT (N2)** | AT3q130 0.0001 | 25.9 ± 10.7 | WT 9.7 ± 4.5 | | *p*=0.065 | |
| | AT3q130 0.001 | 27.2 ± 9.2 | WT 9.7 ± 4.5 | | *p*=0.040 | |
| | AT3q130 0.01 | 22.4 ± 5.0 | WT 9.7 ± 4.5 | | ***p*=0.216** | |
| | AT3q130 0.1 | 22.1 ± 7.0 | WT 9.7 ± 4.5 | | ***p*=0.193** | |
| | AT3q130 1 | 28.6 ± 7.4 | WT 9.7 ± 4.5 | | *p*=0.024 | |
| | AT3q130 10 | 32.2 ± 8.9 | WT 9.7 ± 4.5 | | *p*=0.005 | |
| | AT3q130 50 | 30.4 ± 12.4 | WT 9.7 ± 4.5 | | *p*=0.011 | |
| | AT3q130 100 | 32.5 ± 7.9 | WT 9.7 ± 4.5 | | *p*=0.004 | |

**Table S7** – Descriptive statistics regarding the motility defects of AT3q130 animals upon post-symptomatic CLR01 treatment. Comparisons between groups was carried out using an independent-sample t-test.

| **Strain** | **Treatment** | **Concentration (µM)** | **Mean locomotion defective ± SD** | **Group comparison (Mean locomotion defective ± SD)** | **p-value** |
|---|---|---|---|---|---|
| **AT3q130 (AM685)** | Day 4 (Before treatment) | 0 (vehicle) | 45.1 ± 2.3 | WT 14.2 ± 1.3 | ***p*<0.001** |
| | Day 6 | 0 (vehicle) | 53.9 ± 1.3 | WT 24.4 ± 4.5 | ***p*<0.001** |
| | | 0.1 | 44.4 ± 3.9 | AT3q130 vehicle 53.9 ± 1.3 | ***p*=0.004** |
| | | 0.1 | 44.4 ± 3.9 | WT 24.4 ± 4.5 | ***p*=0.001** |
| | Day 8 | 0 (vehicle) | 55.3 ± 9.1 | WT 33.4 ± 6.4 | ***p*=0.008** |
| | | 0.1 | 41.0 ± 12.9 | AT3q130 vehicle 55.3 ± 9.1 | *p*=0.121 |
| | | 0.1 | 41.0 ± 12.9 | WT 33.4 ± 6.4 | *p*=0.334 |

**Table S8**- Statistical report of CLR01 treatment in *C. elegans*. Effect size calculated using Uanhoro, J. O. (2017), available online at: https://effect-size-calculator.herokuapp.com/.

| **Figure** | | **Statistical report** | **Sample size (n)** |
|---|---|---|---|
| Fig. 5a (chronic treatment (motility)) | | F (8,29) = 3.500; p = 0.006; $\eta p^2 = 0.491$ | 4-5 |
| Fig. 5b (WT chronic treatment (motility)) | | t (6) = 0.103; p = 0.921; g = 0.06 | 4 |
| Fig. 5c (post-symptomatic treatment (motility)) | Day 4 | t (6) = 23.313; p < 0.001; g = 14.13 | 4 |
| | Day 6 | t (6) = 4.648; p = 0.004; g = 2.86 | 4 |
| | Day 8 | t (6) = 1.808; p = 0.121; g = 1.11 | 4 |
| Fig. 5d | Number of aggregates/Total area | t (59) = 0.307; *p* = 0.760 | 31-30 |
| | Area of aggregates | t (59) = 0.130; *p* = 0.897 | 31-30 |
| Fig. 5e | Number of aggregates/Total area | t (60) = 0.883; *p* = 0.384 | 23-39 |
| | Area of aggregates | t (58) = 1.235; *p* = 0.222 | 22-38 |
| Fig. 5f | Number of aggregates/Total area | t (26) = 0.038; *p* = 0.970 | 16-12 |
| | Area of aggregates | t (25) = 6.710; ***p* <0.001** | 15-12 |
| Fig. 5g | Number of aggregates/Total area | t (53) = 0.629; *p* = 0.532 | 27-28 |
| | Area of aggregates | t (53) = 1.495; *p* = 0.141 | 27-28 |

**Table S9.** Reports of all statistical analyses performed in the mouse

<table>
<tr><th>Figure</th><th>Statistical report</th><th>Sample size</th></tr>
<tr><td>CAG repeat</td><td>U = 93.05, p=0.536</td><td>WT VEH, n=19<br>WT CLR01, n=14<br>Q135 VEH, n=19<br>Q135 CLR01, n=13</td></tr>
<tr><td rowspan="3">Body weight</td><td>Group: $F_{(3, 33)}=12.94$, $p < 0.001$, $\eta^2_p = 0.546$</td><td rowspan="3">WT VEH, n=19<br>WT CLR01, n=14<br>Q135 VEH, n=19<br>Q135 CLR01, n=13</td></tr>
<tr><td>Genotype: $F_{(1, 33)}=36.67$, $p < 0.001$, $\eta^2_p = 0.512$</td></tr>
<tr><td>Treatment: $F_{(1, 33)}=0.270$, $p = 0.607$, $\eta^2_p = 0.008$</td></tr>
<tr><td rowspan="3">Motor swimming test</td><td>Group: $F_{(3, 48)}=23.36$, $p < 0.001$, $\eta^2_p = 0.594$</td><td rowspan="3">WT VEH, n=19<br>WT CLR01, n=14<br>Q135 VEH, n=19<br>Q135 CLR01, n=13</td></tr>
<tr><td>Genotype: $F_{(1, 48)}=43.97$, $p < 0.001$, $\eta^2_p = 0.478$</td></tr>
<tr><td>Treatment: $F_{(1, 48)}=7.72$, $p = 0.008$, $\eta^2_p = 0.138$</td></tr>
<tr><td rowspan="3">Beam Square 12mm</td><td>Group: $F_{(3, 45)}=16.43$, $p < 0.001$, $\eta^2_p = 0.523$</td><td rowspan="3">WT VEH, n=19<br>WT CLR01, n=14<br>Q135 VEH, n=19<br>Q135 CLR01, n=13</td></tr>
<tr><td>Genotype: $F_{(1, 45)}=44.74$, $p < 0.001$, $\eta^2_p = 0.499$</td></tr>
<tr><td>Treatment: $F_{(1, 45)}=3.16$, $p = 0.082$, $\eta^2_p = 0.066$</td></tr>
<tr><td rowspan="3">Gait quality</td><td>Group: $H_{(3)}=59.89$ $p<0.001$</td><td rowspan="3">WT VEH, n=19<br>WT CLR01, n=14<br>Q135 VEH, n=19<br>Q135 CLR01, n=13</td></tr>
<tr><td>Genotype: $H_{(1)}=48.46$; $p<0.001$</td></tr>
<tr><td>Treatment: $H_{(1)}=6.83$; $p=0.009$</td></tr>
<tr><td>Aggregation<br>(Neuronal inclusions)<br>in the SCC</td><td>$t_{(4,5)}= 6.748$, $p=0.165$; $d = 0.832$</td><td>5 mice/group (4-7 slices/mouse)</td></tr>
<tr><td>Neuropathology:<br>Pyknotic cells in the<br>SCC</td><td>Group: $F_{(2,14)}=16.011$, $p<0.001$; $\eta^2_p = 0.696$</td><td>WT VEH, n=6; Q135 VEH, n=6;<br>Q135 CLR01, n=5(6 slices/mouse)</td></tr>
<tr><td>Neuropathology:<br>Motor neurons in the<br>SCC</td><td>Group: $F_{(3,16)}=10.397$, $p<0.001$; $\eta^2_p = 0.661$</td><td>WT VEH, n=6; WT CLR01, n=5;<br>Q135 VEH, n=5; Q135 CLR01,<br>n=5(4-6 slices/mouse)</td></tr>
</table>
